# Structure of the Tilapia lake virus nucleoprotein bound to RNA

**DOI:** 10.1101/2024.11.26.625387

**Authors:** Benoît Arragain, Martin Pelosse, Karine Huard, Stephen Cusack

## Abstract

Tilapia Lake virus (TiLV) belongs to the *Amnoonviridae* family within the *Articulavirales* order of segmented negative-strand RNA viruses and is highly diverged from more familiar orthomyxoviruses, such as influenza. The viral nucleoprotein (NP), a key component of the replication machinery, packages the viral genome into protective ribonucleoprotein particles. Here we describe the electron cryo-microscopy (cryo-EM) structure of TiLV-NP bound to RNA within *in vitro* reconstituted, small ring-like, pseudo-symmetrical oligomers. Although TiLV-NP is considerably smaller than its influenza counterpart and unrelated in sequence, it maintains the same topology and domain organisation. This comprises a head and body domain between which is a negatively-charged groove where single-stranded RNA binds. In addition, an oligomerisation loop inserts into a hydrophobic pocket in the neighbouring NP, the flexible hinges of which allow considerable variability orientation of neighbouring NPs. Focussed cryo-EM maps (up to 2.9 Å resolution) unambiguously define the 5′ to 3′ direction of the bound RNA, confirmed by double stranded, A-form RNA regions that extrude out from some of the NP-NP interfaces. This is the first description of how RNA binds to an articulaviral NP and superposition with orthomyxoviral NPs suggest that the mode of RNA binding is likely conserved across the *Articulavirales* order.

## INTRODUCTION

Tilapia Lake virus (TiLV), a recently discovered pathogen of tilapia fish (1), belongs to the *Amnoonviridae* family (2) within the *Articulavirales* order of segmented negative-strand RNA viruses (sNSV). It has 10 single-stranded RNA genome segments, each with conserved, quasi-complimentary 3’ and 5’ ends, a characteristic of sNSV (3). The genome segments encode putative viral proteins whose sequences show no homology to any other protein apart from that of segment 1, which has the conserved motifs characteristic of an orthomyxovirus PB1-like polymerase core subunit (3). A recent structural analysis showed that the complete, functional TiLV RNA-dependent RNA polymerase is a heterotrimer comprising the proteins from segments 1-3 (4). Despite being 40% smaller in size and totally diverged in sequence, TiLV polymerase has an architecture similar to influenza polymerase and contains minimised versions of most of the corresponding domains. Furthermore, TiLV polymerase binds the conserved 3’ and 5’ viral or complementary RNA (vRNA or cRNA) ends in a similar fashion to the influenza viral promoter, with a 5’ hook and distal duplex region, in either mode A (with the 3’ end in the polymerase active site) or mode B (with the 3’ end in the secondary binding site). Finally, similar to influenza polymerase, TiLV polymerase is active for RNA synthesis in vitro and can take up distinct ‘transcriptase’ and ‘replicase’ conformations (4).

The other key component of the sNSV replication machinery is the viral nucleoprotein (NP). Multiple copies of NP package the vRNA or cRNA into ribonucleoprotein particles (RNPs), which are the functional templates for transcription and replication. Orthomyxovirus RNPs are flexible rods in which NP-bound RNA is arranged in two antiparallel strands with an overall super-helical twist, with an RNA loop at one end and the polymerase bound to the 3’ and 5’ (anti-)genomic extremities at the other end (5). The RNPs are necessarily flexible and dynamic, since NPs have to be successively stripped off the template and then replaced during RNA synthesis (6,7). Furthermore, encapsidation of nascent genome replicates is a complex co-replicational process. It is thought to be mediated by the replication complex, an asymmetric dimer, which for influenza, is composed of two viral polymerases bridged by host factor ANP32 (8-10). ANP32 has been proposed to promote co-replicational encapsidation of the elongating replicate within the replication complex by recruitment of successive NPs, thus engendering a progeny RNP (11).

The high resolution structure of the apo-NP is known for several orthomyxoviruses: influenza A (12-15), influenza B (16), influenza D(17), Thogoto (THOV) (18), infectious salmon anaemia (ISAV) (19). Orthomyxovirus NPs exhibit a common fold and NP-NP interaction mechanism, but the inter-subunit flexibility has so far prevented determination of a high-resolution structure of the native influenza or THOV RNP (7,18,20,21). Indeed, the handedness of the RNP helix remains controversial and the mode and direction of binding of the RNA to NP is unknown.

For TiLV, lack of sequence homology precluded straightforward identification of the segment encoding a putative nucleoprotein. However, a previous study has provided convincing experimental evidence that segment 4 encodes the TiLV nucleoprotein (TiLV-NP) (22). The protein was shown to bind in multiple copies to RNA in infected cells and RNP-like particles were immuno-purified from infected cells and virions.

Here we express in insect cells and purify TiLV-NP and biophysically characterise its oligomeric states in apo form and in complex with a 40-mer RNA. By electron cryo-microscopy (cryo-EM), we determine its structure bound to RNA, in the context of small pseudo-symmetrical tetramers, pentamers and hexamers. Despite flexibility within the particles, the resolution is sufficient to unambiguously define the TiLV-NP structure and the mode and directionality of RNA binding. Although smaller than the influenza NP (354 residues compared to 498 for FluA/NP) and with no sequence homology, the topology of the TiLV-NP fold resembles that of orthomyxoviruses. The implications for RNP structure and dynamics are discussed.

## MATERIAL AND METHODS

### Cloning, expression and purification of the TiLV nucleoprotein

As previously described (4) the 10 TiLV open reading frames, codon-optimized for insect cell expression (Genscript), were subcloned into multiple pFastBac Dual vectors using EcoRI and SpeI. The TiLV segment 4, encoding the TiLV nucleoprotein was amplified and inserted into a pACE plasmid with an N-terminal deca-poly-histidine tag followed by a Tobacco Etch Virus protease cleavage site using a combination of PCR and Gibson Assembly (NEB).

The EMBacY bacmid containing the TiLV nucleoprotein gene was prepared using the Bac-to-Bac method (Invitrogen) and subsequently used for insect cell expression. For large-scale expression, Trichoplusia ni High 5 cells (ThermoFisher) at a concentration of 0.8–1 × 10^6^ cells/mL were infected by adding 1% of the virus. Expression was stopped 72 to 96 hours after the day of proliferation arrest, and the cells were harvested by centrifugation (1000 g, 20 min, 4 °C). The cells were disrupted by sonication for 3 minutes (10 s ON, 20 s OFF, 40% amplitude) on ice in lysis buffer (50 mM HEPES pH 8, 500 mM NaCl, 20 mM imidazole, 0.5 mM TCEP, and 5% glycerol) with cOmplete EDTA-free Protease Inhibitor Cocktail (Roche). After lysate centrifugation at 48,000 g for 45 minutes at 4 °C, the soluble fraction was loaded on a HisTrap HP ion affinity chromatography column (Cytiva). Bound proteins were subjected to two sequential wash steps using (i) the lysis buffer supplemented with 1 M NaCl and (ii) the lysis buffer supplemented with 50 mM imidazole. Bound proteins were eluted gradually from 50 mM imidazole to 500 mM imidazole over 15 column volume.

TiLV nucleoprotein fractions were pooled. Tobacco Etch Virus protease was added for His-tag cleavage (1:50 w/w ratio), and the protein mixture was dialyzed overnight at 4 °C in a heparin-loading buffer (50 mM HEPES pH 8, 250 mM NaCl, 0.5 mM TCEP, 5% glycerol). Proteins were loaded on a HiTrap Heparin HP column (Cytiva), washed using the heparin-loading buffer, and eluted gradually from 250 mM NaCl to 1 M NaCl, over 15 column volume, using 50 mM HEPES pH 8, 1 M NaCl, 2 mM TCEP, 5% glycerol. The nucleic acid-free TiLV nucleoprotein fractions (ratio A260/280 = 0.56) were dialyzed overnight at 4 °C in a final buffer (50 mM HEPES pH 8, 300 mM NaCl, 0.5 mM TCEP, 5% glycerol), concentrated using Amicon Ultra (10 kDa cut-off), flash-frozen in liquid nitrogen, and stored at −80 °C for further use.

### Size-exclusion chromatography with multi-angle light scattering

SEC-MALS experiments were performed on an OMNISEC system (Malvern) using a Superdex 200 Increase 10/300 GL column (Cytiva). The column was equilibrated with a buffer containing 50 mM HEPES pH 8, 150 mM NaCl, 2 mM TCEP. The system calibration was performed using a 50 µL injection of Bovine Serum Albumin (BSA) at 2 mg/ml to identify the monomeric and dimeric populations. Following calibration, 50 µL injections of apo TiLV-NP at concentrations of 5 mg/ml, 3 mg/ml, and 3 mg/ml in the presence of the 40-mer vRNA loop (5′-pGCA AAU CUU UCU CAC GUC CUG ACU UGU GAG UAA AAU UUG G −3′) (1 NP : 0.5 RNA molar ratio) were performed. Static light scattering detection was conducted using the OMNISEC system′s RALS and LALS detectors (Malvern) with a laser emitting at 660 nm. The weight-average molar masses were calculated using the OMNISEC v5.10 software (Malvern).

### Mass photometry analysis

Mass photometry measurements were performed on a OneMP mass photometer (Refeyn). Coverslips (No. 1.5H, 24 × 50 mm, VWR) were washed with water and isopropanol before being used as a support for silicone gaskets (CultureWellTM 423 Reusable Gaskets, Grace Bio-labs). Contrast/mass calibration was realised using native marker (Native Marker unstained protein 426 standard, LC0725, Life Technologies) with a small field of view and monitored during 60 s using the AcquireMP software (Refeyn). For each condition, 18 µl of buffer (50 mM HEPES pH 8, 150 mM NaCl, 2 mM TCEP) were used to find the focus. 2 µl of sample were added to reach a final TiLV-NP concentration of 25 nM. Movies of 60 s were recorded, processed and mass estimation was determined automatically using the DiscoverMP software (Refeyn).

### Sample preparation, cryo-EM grid freezing and data collection

For sample preparation, 100 µM apo TiLV-NP were incubated with the 40-mer vRNA loop (5’-pGCA AAU CUU UCU CAC GUC CUG AUU UGU GAG UAA AAU UUG G -3’) (1:0.5 molar ratio) for 1h at 4°C in a SEC buffer containing 50 mM HEPES pH 8, 150 mM NaCl, 2 mM TCEP. The resulting mixture was centrifuged 5 min at 11,000 g prior to injection onto a Superdex 200 Increase 3.2/300 column (Cytiva). To enrich for larger RNA-bound oligomers (≥ tetramers), two adjacent left-side fractions of the main SEC peak were collected and used to optimize the TiLV-NP-RNA concentration used for cryo-EM grid preparation. For cryo-EM grid preparation, 1.5 µl of sample was applied on each side of plasma cleaned (Fischione 1070 Plasma Cleaner: 1 min 30, 80% oxygen, 20% argon) grids (UltrAufoil 1.2/1.3, Au 300). Excess solution was blotted for 3 sec, blot force -2, at 100% humidity and 4°C with a Vitrobot Mark IV (ThermoFisher) before plunge freezing in liquid ethane.

Two automated data collections were performed on two different grids using a TEM Krios (ThermoFisher) operated at 300 kV equipped with a K3 direct electron detector camera (Gatan) using SerialEM (23). Coma and astigmatism correction were performed on a carbon grid. Movies of 40 frames were recorded in counting mode at a ×105,000 magnification, giving a pixel size of 0.822 Å, with defocus ranging from −0.8 to −2.0 μm. Total exposure dose was ∼40 e−/Å^2^ or ∼50 e−/Å^2^.

### Image processing

For the two collected datasets, the image processing followed a similar workflow carried out in CryoSPARC v4.3 and v4.5 (24). Movie drift correction was performed using all frames, along with gain reference and camera defect corrections. CTF parameters were determined using “Patch CTF estimation”. Realigned micrographs were then inspected and low-quality images were manually discarded. RNA bound TiLV-NP oligomers were automatically picked using a circular blob with diameters ranging from 80 to 140 Å. Particles were extracted, 2D classified, and subjected to an “Ab-initio reconstruction” job to generate multiple initial 3D reconstructions. The best 3D models were used for one round of heterogeneous refinement and the resulting 3D classes subjected to a non-uniform 3D refinement job. The resulting maps were used to prepare 2D templates and particles picked using the template picker job. Resulting particles were combined, with duplicates removed, and extracted using a box size of 340 × 340 pixels^2^. Successive 2D classifications were used to eliminate particles displaying poor structural features. A second heterogeneous refinement was performed using tetramers (pseudo-C2 and pseudo-C4), pentamer (pseudo-C5) and hexamer (pseudo-C6) maps as initial 3D models. Particles belonging to a specific oligomer were re-centered and re-extracted, 2D classified to ensure that there is the least misclassification, and subjected to non-uniform 3D refinement, which was then used for reference-based motion correction (25). Motion-corrected particles were subjected to a last non-uniform 3D refinement and resulted in four final maps, one for each RNA-bound TiLV-NP oligomers (pseudo-C2/-C4/-C5/-C6).

The inherent flexibility between each TiLV-NP prevented application of a defined symmetry. Based on the overall complete C1 maps (pseudo-C2/-C4/-C5/-C6), symmetry expansion was performed using C2/C4/C5 and C6 transformation, respectively. To better account for NP-NP interactions, two TiLV-NPs were kept while the remaining regions were subtracted. The resulting particles were used for local 3D refinement. Finally, 3D classification without alignments were performed to isolate the most homogeneous subset of particles, itself subjected to a final local refinement, resulting in four final maps enclosing 2 TiLV-NPs for each RNA-bound TiLV-NP oligomers (pseudo-C2/-C4/-C5/-C6).

Post-processing was performed in CryoSPARC using an automatically determined B-factor. For each final map, reported global resolution is based on the FSC 0.143 cut-off criteria. Local resolution variations were estimated in CryoSPARC.

For detailed image processing information, please refer to Supplementary Figures 1 and 2. For assessing cryo-EM map quality, please refer to Supplementary Figures 3, 4 and 8.

### Model building and refinement

Model building of the full-length TiLV nucleoprotein and associated RNA (deposited with generic purine or pyrimidine bases) was performed de novo in COOT (26) using the map corresponding to two NPs, from the RNA-bound TiLV-NP pentamer (pseudo-C5), at 2.9 Å resolution. Models were refined using Phenix real-space refinement (27) with Ramachandran restraints. The TiLV-NP model was duplicated and rigidly fitted into the density for each oligomer. The oligomerization loop hinges, the RNA and other flexible loops were then manually adjusted into the cryo-EM density, followed by one round of refinement. For the RNA, exact register of sequence could not be determined and purines or pyrimidines were assigned depending on apparent base size, whilst maintaining Watson-Crick base-pairing in the double-stranded regions. In the deposited PDBs, generic purine (P5P) or pyrimidine (Y5P) nucleotides were substituted. The CCmask is relatively low for the pseudo-C4 and pseudo-C6 structures, attributable to poorer density for 2 and 3 NPs respectively. Atomic model validation (Supplementary Tables 1-4) was performed using MolProbity (28) as implemented in Phenix. Model resolution according to the cryo-EM map was estimated at the 0.5 FSC cutoff. Electrostatic potential was calculated using PDB2PQR and APBS (29). Figures were generated using ChimeraX (30).

## RESULTS

### Expression, purification and biophysical characterization of apo TiLV-NP and RNA-bound TiLV-NP oligomers

TiLV-NP is a 38.4 kDa protein composed of 354 amino acids and encoded on the TiLV segment 4 (22). TiLV-NP was expressed in insect cells and purified in its apo form, as confirmed by an A260/280 ratio of 0.56 indicating the absence of nucleic acids (Fig. 1A, B; see Material and Methods). Standard size exclusion chromatography (SEC) revealed that apo TiLV-NP elutes as a broad peak, suggesting the presence of multiple oligomeric states in solution, as seen for instance for FluA/NP (13). Further analysis using SEC coupled with multi-angle light scattering (SEC-MALS) showed that apo TiLV-NP predominantly form trimers and tetramers in roughly equal proportions, regardless of protein concentration. At 5 mg/ml, ∼53% tetramers and ∼46% of trimers were detected, while at 3 mg/ml, it formed ∼45% tetramers and ∼54% trimers (Fig. 1C). The experimental molecular weight of the apo TiLV-NP tetramers (5 mg/ml: 161.6 kDa; 3 mg/ml: 156.1 kDa) is within 5% of the theoretical value (153.6 kDa). However, the apo TiLV-NP trimeric form showed a larger discrepancy with the theoretical molecular weight (115.2 kDa) corresponding to only 86% of the experimental value (134.1 and 133 kDa), perhaps indicating some flexibility or conformational changes (Fig. 1C). Complementary mass photometry analysis, performed at nM protein concentration (versus µM for SEC), yielded similar results. Two main peaks are observed at 109 kDa (trimers, 33%) and 150 kDa (tetramers, 53%). In addition, a third less-resolved peak, corresponding to a pentameric form was detected, albeit in lower amounts (14%), a species not detected in SEC-MALS (Fig. 1D).

**Figure 1.**
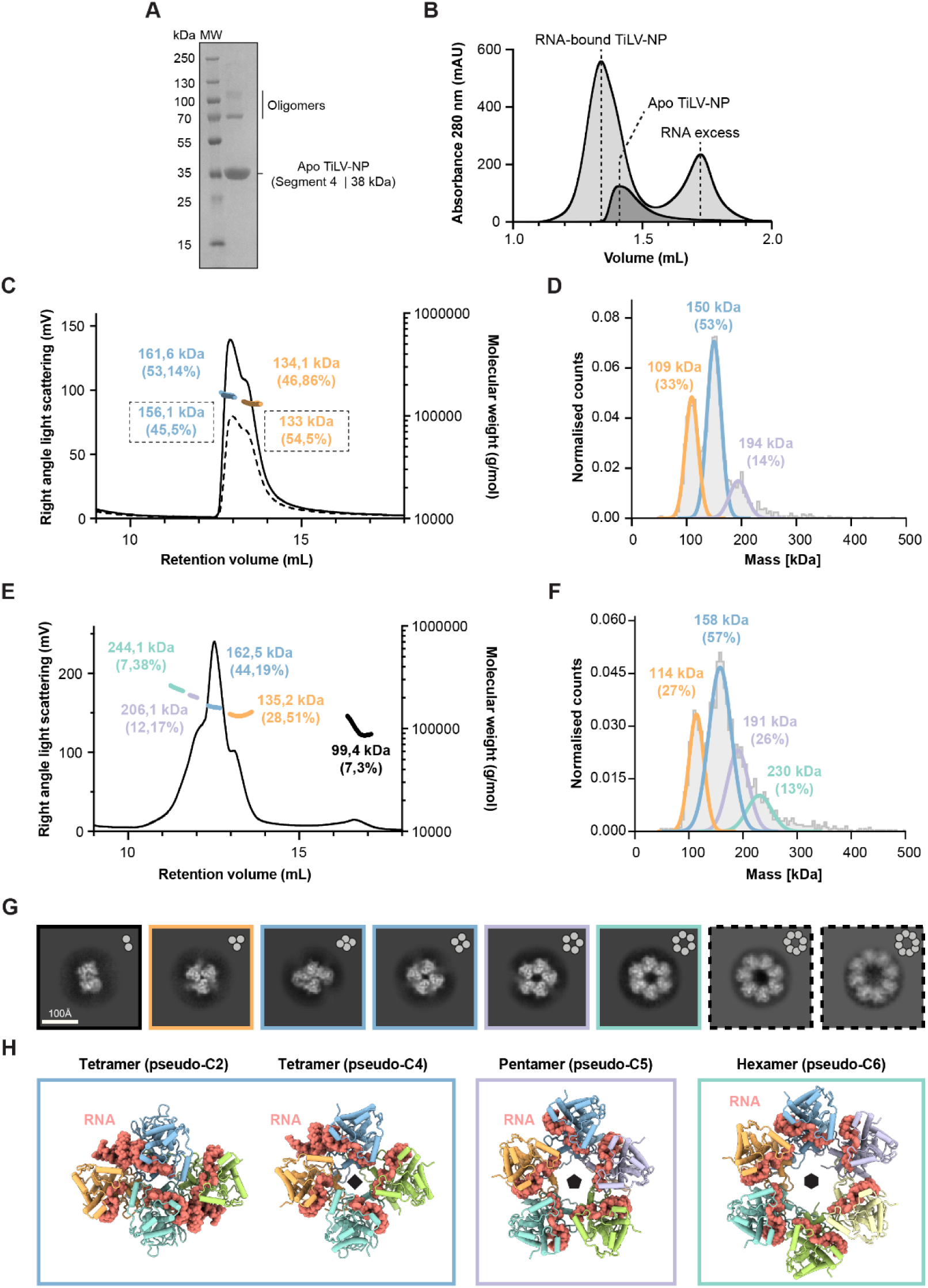
TiLV-NP biochemical, biophysical, and structural characterizations. A. SDS-PAGE analysis of purified apo TiLV-NP. The molecular ladder (MW) is shown on the left side of the gel. Apo TiLV-NP is indicated on the right side of the gel (segment 4; 38.4 kDa). Despite denaturing conditions, apo TiLV-NP oligomers remain visible and indicated on the right side of the gel. B. Size exclusion chromatography (SEC) profiles of RNA-bound TiLV-NP (light grey) and apo TiLV-NP (dark grey). The relative absorbance at 280 nm (mAU) is on the y-axis. The elution volume (mL) is on the x-axis, graduated every 100 µL. C. SEC coupled with multi-angle light scattering (MALS) of apo TiLV-NP injected at 5 mg/mL (black solid line) and 3 mg/mL (black dotted line). The right angle light scattering (mV) is on the left y-axis, the molecular weight (g/mol) on the right y-axis. The elution volume (mL) is on the x-axis, graduated every 1 mL. The molecular weights of apo TiLV-NP tetramers and trimers are coloured in blue and orange, outlined for the 3 mg/mL run. D. Mass photometry analysis of apo TiLV-NP. The curves and molecular weights for apo TiLV-NP pentamers, tetramers and trimers are coloured mauve, blue and orange, respectively. E. SEC coupled with MALS of RNA-bound TiLV-NP injected at 3 mg/mL (black solid line). The right-angle light scattering (mV) is on the left y-axis, the molecular weight (g/mol) on the right y-axis. The elution volume (mL) is on the x-axis, graduated every 1 mL. The molecular weights determined for RNA-bound TiLV-NP hexamers, pentamers, tetramers, trimers and dimers are respectively coloured green, mauve, blue, orange and black. F. Mass photometry analysis of RNA-bound TiLV-NP. The curves and molecular weights for apo TiLV-NP hexamers, pentamers, tetramers and trimers are coloured green, mauve, blue, and orange, respectively. TiLV-NP dimers are not detected. G. Cryo-EM 2D class averages of RNA-bound TiLV-NP oligomers. Dimers and trimers are outlined in black and orange, respectively. Tetramers are present in two distinct conformations and outlined in blue. Pentamers and hexamers are outlined in mauve and green, respectively. Heptamers and octamers, which are not detected in SEC-MALS or mass photometry, are outlined with black dotted lines. H. Cartoon representation of the different RNA-bound TiLV-NP structures solved by cryo-EM. Each TiLV-NP is coloured differently (orange, blue, mauve, yellow, green and cyan). The RNA is displayed as spheres, coloured in salmon.

To reconstitute RNA bound TiLV-NP oligomers, apo TiLV-NP was incubated with the previously described 40-mer vRNA loop (vRNA loop, 5’-pGCA AAU CUU UCU CAC GUC CUG AUU UGU GAG UAA AAU UUG G -3’), which corresponds to the joining of the partially complementary first (5′) and last (3′) 20 nucleotides of the ends of TiLV vRNA segment 9 (4). Comparative SEC analysis indicated a shift and peak broadening in the presence of RNA compared to the apo TiLV-NP, suggesting an enhanced oligomerization (Fig. 1B). This was further confirmed by SEC-MALS, which showed that RNA-bound TiLV-NP forms oligomers ranging from dimers to hexamers (Fig. 1E). Trimers and tetramers were still the most abundant, as for apo TiLV-NP, with ∼44% and ∼29% of the total population, respectively. The pentameric form accounted for ∼12%, while the hexameric and dimeric forms are present at ∼7% each. The experimental molecular weights matched closely with theoretical masses if one assumes that the TiLV-NP oligomers are bound to at least one copy of the vRNA loop (12.8 kDa): 244.1 kDa (experimental) versus 243.2 (theoretical) for hexamers (99.6%), 206.1 kDa versus 204.8 for pentamers (99.4%), 162.5 kDa versus 166.4 for tetramers (102.4%), 135.2 kDa for 128 for trimers (94.6%), and 99.4 kDa for 89.6 for dimers (90.1%) (Fig. 1E). The presence of dimers, not detected in the apo TiLV-NP SEC-MALS analysis, suggests that apo TiLV-NP trimers and tetramers could possibly dissociate to accommodate RNA binding and subsequently form higher-order oligomers. Complimentary mass photometry analysis validated these findings with an increased presence of pentamers (26%) and the appearance of hexamers (13%), although peak resolution remains limited. The dimeric population is not observed (Fig. 1F).

Taken together, these biophysical results show that apo TiLV-NP exist predominantly in trimeric and tetrameric forms, whilst RNA binding drives the oligomerization of TiLV-NP towards higher order oligomers, principally pentamers and hexamers.

### Cryo-EM structure of the RNA-bound TiLV-NP and comparison with the apo influenza A/NP

To structurally characterize RNA-bound TiLV-NP oligomers, a mixture of vRNA loop and TiLV-NP was incubated and loaded on SEC (see Materials and Methods). To enrich for high-order oligomers (≥ pentamers), different early fractions of the SEC peak were deposited on grids, vitrified, and imaged by electron cryo-microscopy (cryo-EM) (Supplementary Fig. 1, 2). Initial 2D classifications revealed a diverse population of RNA-bound TiLV-NP oligomers, with pseudo-symmetry from C2 to C8, with octamers being the highest observed oligomeric state (Fig. 1G). No pseudo-helicoidal assemblies were detected under these in vitro conditions. Following cryo-EM data collection and image processing, multiple pseudo-symmetrical TiLV-NP oligomer structures bound to the vRNA loop were resolved, with overall resolution ranging from 2.9 Å to 3.7 Å (Fig. 1H; Supplementary Tables 1-4; Supplementary Fig. 1-3). These structures include two distinct tetrameric forms (pseudo-C2 and pseudo-C4), one pentameric (pseudo-C5) and one hexameric (pseudo-C6) form. The pseudo-C2 and pseudo-C5 oligomers appeared the most stable with uniform density according to local resolution, while the pseudo-C4 and pseudo-C6 structures displayed poorer density for 2 and 3 NPs, respectively (Supplementary Fig. 1-3). To account for this variability, symmetry expansion, particle subtraction, 3D classification without alignments, and local refinements were applied during cryo-EM image processing. This enabled isolation of the most homogeneous sub-oligomers, corresponding to two NPs within each oligomer, and deciphering of their respective NP-NP interactions (Supplementary Fig. 1-3). The determination of higher oligomeric state structures (≥ pseudo-C7) was hindered by the absence of side views and a low number of particles. Conversely, lower oligomeric state structures (≤ pseudo-C3) lacked sufficient signal-to-noise ratio, primarily due to the ice thickness relative to the particle size, impeding accurate particle picking, sorting, and angle estimation for 3D reconstruction.

TiLV-NP is 30% smaller than the influenza A nucleoprotein (FluA/NP), consisting of 354 amino acids versus 498 for FluA/NP. This size reduction is analogous to that observed for the heterotrimeric TiLV viral polymerase, which is only 60% the size of the influenza A polymerase (4). Nevertheless, TiLV-NP adopts a topologically similar fold to FluA/NP. It is rich in α-helices and adopts a crescent-like shape divided into three subdomains similar to orthomyxovirus NPs (Fig. 2; Supplementary Fig. 4, 5). Of the 354 residues, the first 33 N-terminal amino acids are not visible in any cryo-EM maps, presumably due to flexibility. The TiLV-NP “body” domain comprises residues 34-175, 228-287, 320-350, and contains an extending oligomerization loop (294-307) flanked by flexible hinges (288-293; 308-319). The “head” domain (176-227) is inserted within the body domain and separates two critical regions (Fig. 2A). On one side, along with the body domain, is formed a positively charged RNA-binding groove that can accommodate up to 12 nucleotides (Fig. 2A, B) and on the other side, a hydrophobic groove into which the oligomerization loop (‘tail-loop’ in Flu/NP) of an adjacent TiLV-NP can insert (Fig. 2B). Compared to FluA/NP (Fig. 2C, D), the TiLV-NP body domain is the least reduced, retaining 80% of its size (230 versus 287 residues). In contrast, the TiLV-NP head domain is the most prominently minimized, consisting of only three α-helices and being 38% the size of the FluA/NP counterpart (51 residues versus 136). Finally, the TiLV-NP oligomerization loop and both hinges are also reduced in size and does not extend as much as in FluA/NP (31 versus 40 residues) (compare Fig. 2A, B and C, D).

**Figure 2.**
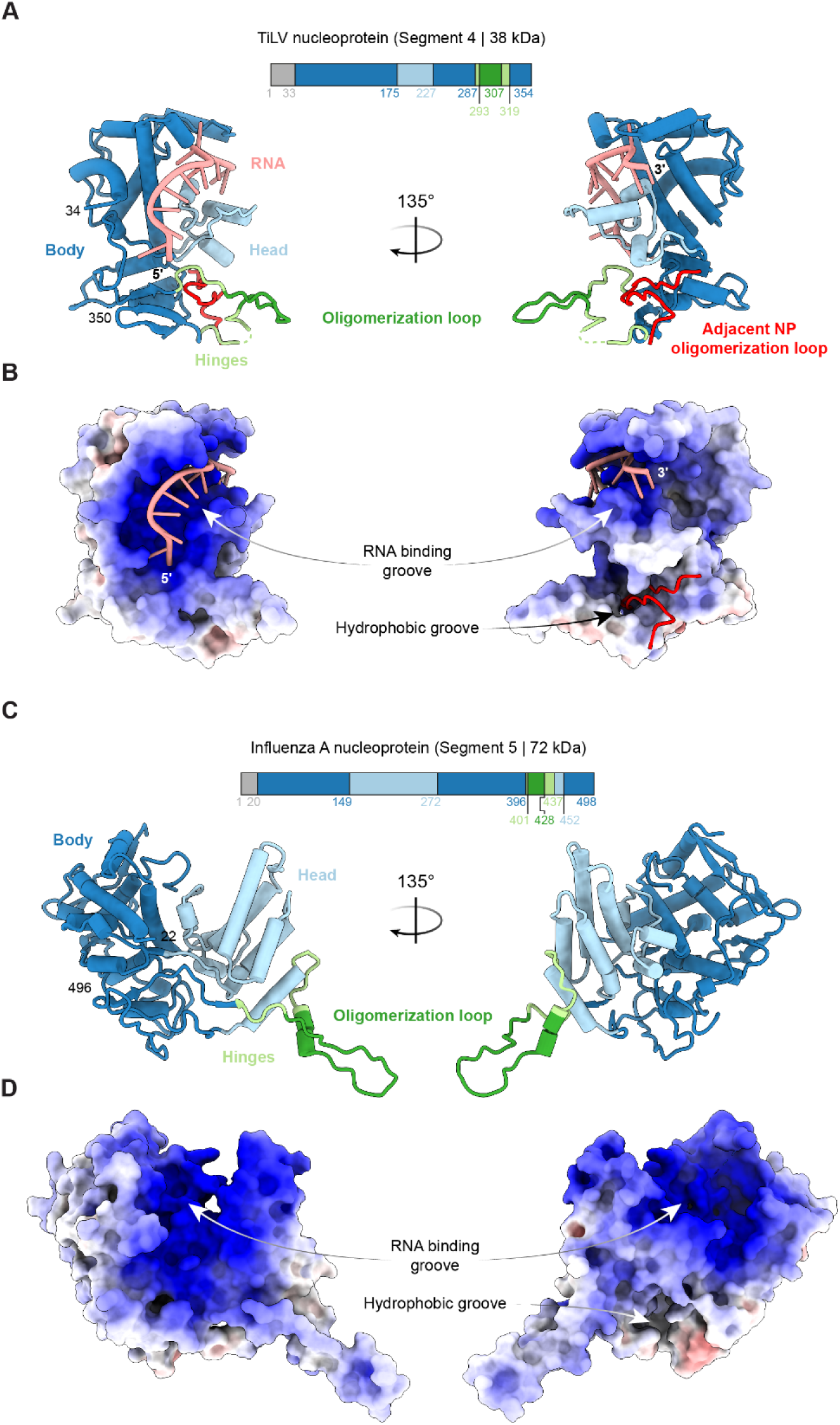
Structure of the RNA-bound TiLV-NP and comparison with FluA/NP. A. Schematic of the TiLV-NP domains and cartoon representation of the RNA-bound TiLV-NP structure. The first N-terminal 33 residues are unseen (grey). The body domain is coloured in dark blue, the head domain in light blue, the hinges in light green, the RNA in salmon, and the oligomerization loop in dark green. The adjacent NP oligomerization loop is coloured in red. B. Electrostatic surface representation of TiLV-NP structure. Dark blue is positively charged regions (+10), white is neutral (0) and red is negative (-10). The RNA binding groove and the hydrophobic groove, in which the adjacent NP oligomerization loop docks, are annotated. C. Schematic of the FluA/NP domains and cartoon representation of the apo FluA/H5N1-NP structure (PDB 2Q06). The first N-terminal 20 residues are unseen (grey). All domains are coloured as for TiLV-NP in panel A. D. Electrostatic surface representation of FluA/NP structure. Dark blue is positively charged regions (+10), white is neutral (0) and red is negative (-10). The RNA binding groove and the hydrophobic groove, in which the adjacent NP oligomerization loop docks, are annotated.

### Overall structure of the RNA-bound TiLV-NP oligomers and NP-NP interactions

The RNA-bound TiLV-NP structure has been determined as part of distinct pseudo-symmetrical oligomers, including two tetrameric forms with pseudo-C2 and pseudo-C4 symmetries (Fig. 1H). The pseudo-C2 tetramer adopts an elongated structure, measuring 116 Å in length, 99 Å in width, and 70 Å in height (Supplementary Fig. 6A). In contrast, the pseudo-C4 tetramer is more compact, with dimensions of 95 Å in length, 102 Å in width, and 62 Å in height (Fig. 1H; Supplementary Fig. 6B). This structural variability is due to distinct RNA binding modes at the NP-NP interfaces. Within the pseudo-C2 tetramer, two A-form RNA double-stranded helices are inserted at the interfaces between opposite pairs of NP, whose single-stranded 5’ and 3’ extensions bind each of the four NPs (Fig. 1H; Supplementary Fig. 6A). For the pseudo-C4 tetramer, the observed RNA is mostly single stranded, with one less well-ordered, double-stranded region (Fig. 1H; Supplementary Fig. 6B). Consequently, the NP-NP interface areas differ. The pseudo-C2 tetramer exhibits mean interface area of 1294 Å^2^ between NPs within one dimer and 994 Å2 at the dimer interface, while the pseudo-C4 tetramer exhibits a more uniform interface of 1313 Å^2^ between NPs. In addition to the two tetramers, pentameric and hexameric oligomers were also resolved, displaying pseudo-C5 and pseudo-C6 symmetries (Fig. 1H; Supplementary Fig. 6C, D). These higher-order oligomers resemble the pseudo-C4 structure but incorporate additional NPs, resulting in circular assemblies with diameters of ∼107 Å for the pentamer and ∼122 Å for the hexamer (Supplementary Fig. 6C, D). The NP-NP interface areas are smaller, averaging 1240 Å² for the pentamer and 1065 Å² for the hexamer. The accommodation of an A-form double-stranded RNA helix at the pseudo-C2 NP interface, as well as the oligomerization into circular pentamers and hexamers, is facilitated by the flexible hinges of TiLV-NP (residues 288-293, 308-319). These allow for slight shifts and rotations between adjacent NPs, whilst maintaining strong NP-NP interactions via the interpenetrating oligomerisation loop (Fig. 3A, B; Supplementary Fig. 6A-D).

**Figure 3.**
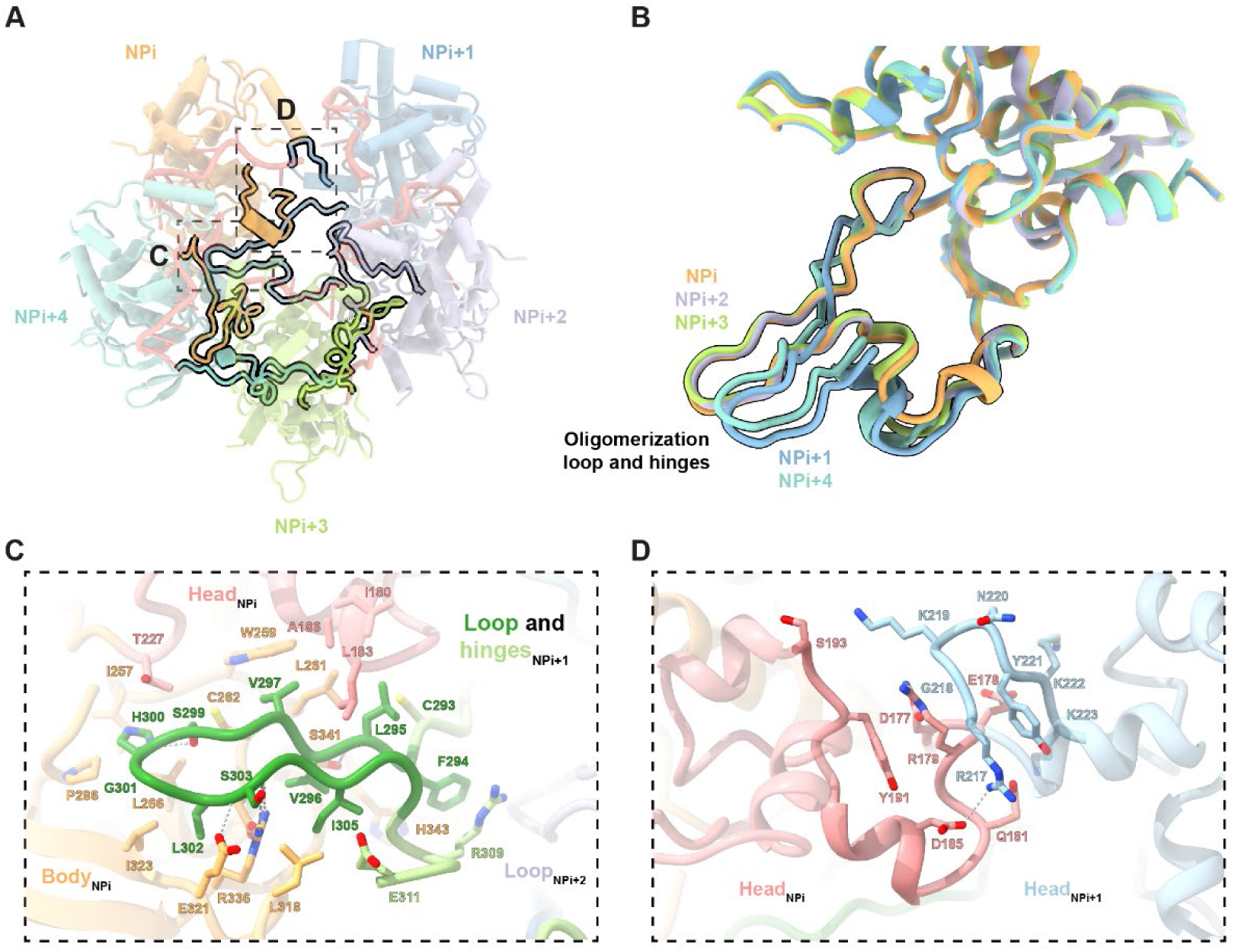
TiLV-NP oligomerization and NP-NP interfaces. A. Cartoon representation of the TiLV-NP pseudo-C5 structure, and highlight of the different NP-NP interface areas. TiLV-NP pseudo-C5 structure, with NPs coloured in orange (NPi), dark blue (NPi+1), mauve (NPi+2), green (NPi+3), cyan (NPi+4), is shown in transparent. The oligomerization loop of each NP and the head-to-head interaction between NPi and NPi+1 are highlighted with black solid lines. Dotted squares indicate close-up views presented in panels C and D. B. Superposition of each NP from the TiLV-NP pseudo-C5 structure. The oligomerization loop and hinges take two main conformations: one shared by NPi, NPi+2 and NPi+3, and another by NPi+1 and NPi+4. NPs are coloured as in A. The oligomerization loop and hinges of each NP are highlighted with black solid lines. C. Close-up view of the interaction between the TiLV-NPi+1 oligomerization loop and the TiLV-NPi body and head domains. TiLV-NPi body is coloured in orange and the head domain in pale red. TiLV-NPi+1 domains are coloured as in Fig. 2A. Interacting residues are shown as sticks, in non-transparent. Ionic and hydrogen bonds are shown as grey dotted lines. D. Close-up view of the interaction between TiLV-NPi+1 oligomerization loop and TiLV-NPi body and head domains. Ionic and hydrogen bonds are shown as grey dotted lines.

TiLV-NP oligomerization is mediated by two main contact areas between adjacent NPs (Fig. 3A). The primary interaction, conserved within all oligomers, is formed by the β-hairpin-like oligomerization loop of TiLV-NPi+1 (residues 293-308 and E311), which inserts into a hydrophobic pocket formed by both the body (residues 336-343, 318-324, 257-266, 286-289) and head (residues 180-186, 227) domains of the adjacent TiLV-NPi (Fig. 3A). Key interactions include the stacking of TiLV-NPi+1 F294, stabilised by R309, with TiLV-NPi H343. The first strand of TiLV-NPi+1 293-CFLVVA is stabilised by TiLV-NPi head (I180, L183, A185 residues) and TiLV-NPi+1 body (L261, W259 and C262 residues) (Fig. 3C). TiLV-NPi+1 tip residues S299 and H300 are inserted into a pocket formed by TiLV-NPi T227, I257 and L266, with TiLV-NPi+1 G301 stabilised by TiLV-NPi I323 (Fig. 3C). The second strand (TiLV-NPi+1 302-LSAI) is stabilised by TiLV-NPi E321, R336, L318, with TiLV-NPi+1 L302 being clamped between TiLV-NPi I323, E321 and R336 (Fig. 3C). TiLV-NPi+1 S303 amine and carboxyl groups are hydrogen-bonded to TiLV-NPi E321 and R336, respectively (Fig. 3C). These interactions account for ∼82% (∼940A^2^) of the total interface area between adjacent TiLV-NPi+1 and TiLV-NPi, making them essential for oligomeric assembly.

The second area is located at the interface between both head domains of TiLV-NPi (191-193, 178-185) and TiLV-NPi+1 (217-223). Notably, TiLV-NPi D185 interacts with TiLV-NPi+1 R217, while TiLV-NPi Y191 stacks with R179, which in turn interact with TiLV-NPi+1 G218 (Fig. 3D). Additionally, TiLV-NPi S193 is close to TiLV-NPi+1 K219, and TiLV-NPi E178 is surrounded by TiLV-NPi+1 residues 220-223. This secondary interaction is less consistent between NPs, suggesting accommodation depending on the NPi-NPi+1 angle. Conversely, positively charged residues within the 211-225 loop of the head domain are also implicated in the formation of the TiLV-NP RNA binding groove and RNA stabilisation, described in more detail below (Fig. 4).

**Figure 4.**
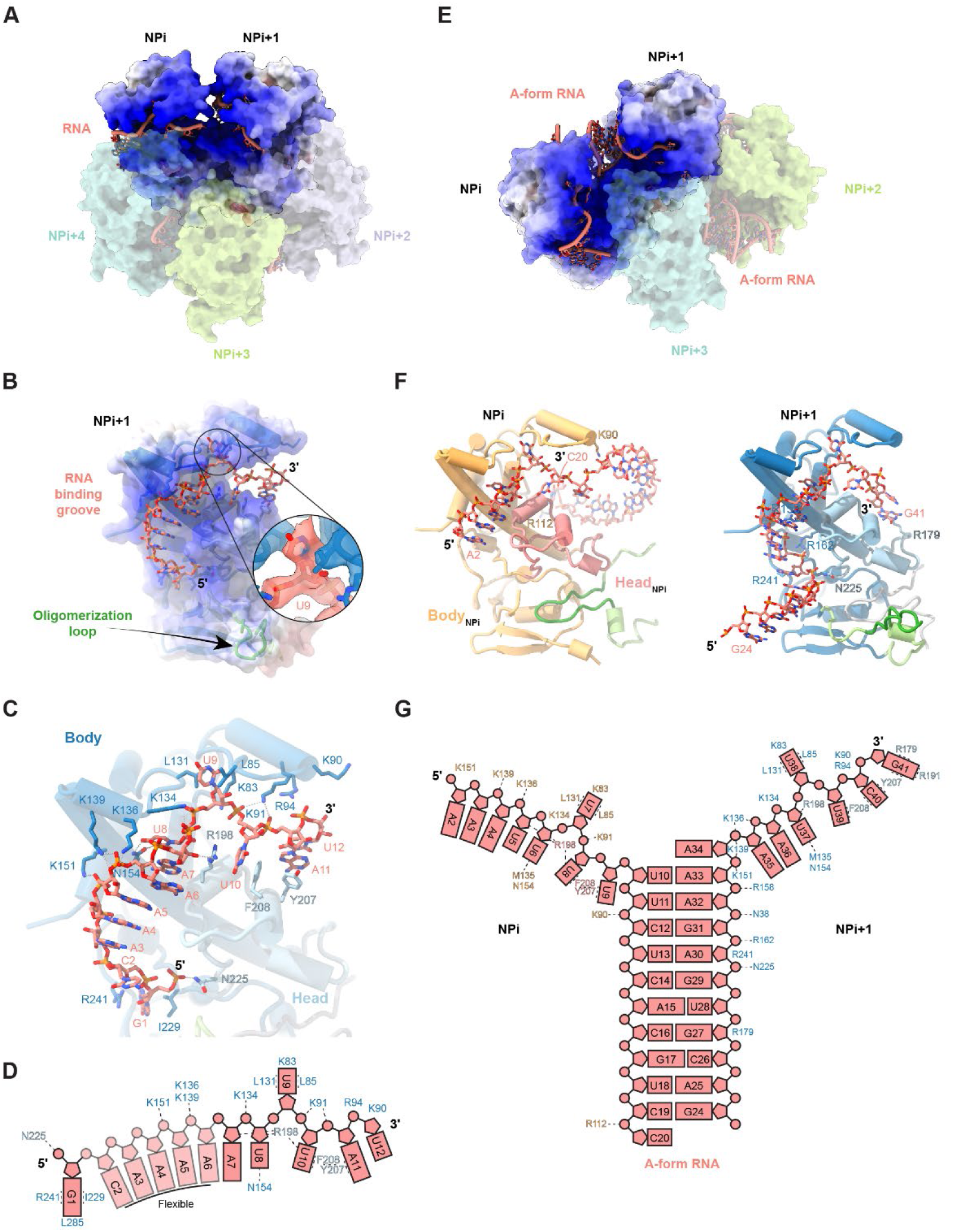
RNA interaction and polarity within the TiLV-NP. A. Surface representation of the RNA-bound TiLV-NP pseudo-C5 structure. Two NPs (NPi, NPi+1) are shown with electrostatic surface representation where dark blue is positively charged regions (+20), white is neutral (0), and red is negative (-20). NPi+2, NPi+3 and NPi+4 are coloured as in Fig. 1A, and shown in transparent. The RNA, coloured in salmon, bind as single strand within each NP. B. Close-up view of the RNA binding groove in TiLV-NPi+1 from the TiLV-NP pseudo-C5 structure. Electrostatic surface representation is shown in transparent with cartoon representation of TiLV-NP structure underneath. The RNA is as sticks. The 5′ and 3′ ends are indicated. The 5′ end is following the oligomerization loop orientation. A close-up view on coulomb potential around U9 unambiguously reveal the RNA polarity. C. Close-up view of the interaction between the RNA and TiLV-NP body and head domains. TiLV-NP domains are coloured as in Fig. 2A. Interacting residues are shown as sticks, in non-transparent. Ionic and hydrogen bonds are shown as grey dotted lines. The extracted RNA density from the local refinement around 2 NPs from the TiLV-NP pseudo-C5 cryo-EM map (2.9 Å) unambiguously show the RNA polarity. D. Schematic summary of the interactions between TiLV-NPi+1 from the local refinement around 2 NPs (pseudo-C5 structure) and the vRNA loop. The RNA is coloured as salmon. Interacting residues are coloured based on the domain they belong to, dark blue for the body domain and light blue for the head domain. Flexible nucleotides are in transparent and indicated. E. Surface representation of the RNA-bound TiLV-NP pseudo-C2 structure. Two NPs (NPi, NPi+1) are shown with electrostatic surface representation, where dark blue is positively charged regions (+20), white is neutral (0) and red is negative (-20). NPi+2 and NPi+3 are coloured as in Fig. 1A, shown in transparent. The RNA bind as single strand within each NP, forming an A-form double-stranded RNA helix at the interface of NPi/NPi+1 and NPi+2/NPi+3. F. Cartoon representation of TiLV-NPi and TiLV-NPi+1 from the TiLV-NP pseudo-C2 structure. Only the RNA strand interacting with each respective NP is shown for clarity. Interacting residues not observed in the TiLV-NP pseudo-C5 structure are indicated. TiLV-NPi domains are coloured as in Fig. 3C. TiLV-NPi+1 domains are coloured as in Fig. 2A. G. Schematic summary of the interactions between TiLV-NPi, TiLV-NPi+1 from the local refinement around 2 NPs (pseudo-C2 structure) and the vRNA loop. The RNA is coloured salmon. Interacting residues are coloured according to the domain they belong as in panel F.

### TiLV-NP RNA binding mode

Within each TiLV-NP oligomer, all NPs bind the RNA as a single strand within a positively charged binding groove formed by both the body and head domains. The RNA is directed towards the central part of each oligomer and is thus protected from solvent, as also observed in NP oligomers from bunyaviruses (Fig. 4A, B; Supplementary Fig. 7). In contrast, the corresponding positively charged groove is solvent exposed in the apo-FluA/H5N1-NP, FluA/H1N1-NP, FluB/Managua/2008-NP, and FluD/Bovine/France-NP oligomers previously characterized using X-ray crystallography (Supplementary Fig. 7).

In the pseudo-C5 oligomers, up to 12 nucleotides can be modelled per NP (Fig. 4A-D). The RNA-binding mode is conserved across all NPs, with ∼930Å^2^ of RNA surface buried within each groove, making the RNA largely inaccessible to the solvent. Only discontinuous RNA density is observed at the NP-NP interfaces suggesting two potential binding modes: either individual vRNA loops interact separately with each NP, or a single vRNA loop may bridge two adjacent NPs, leaving a flexible RNA loop at the NP-NP interface.

In contrast, the pseudo-C2 oligomer structure exhibits a distinct RNA-binding mode (Fig. 4E). Within each of the positively charged grooves of TiLV-NPi and TiLV-NPi+1, 8 nucleotides of RNA are bound, with free 5′ and 3′ ends respectively (Fig. 4F). These two strands then come together at the NP-NP interface to form an A-form RNA double-helix comprising at least 10 base-pairs with a total of around 37 nucleotides visible (Fig. 4G). Two such structures bind to the pseudo-C2 oligomer (Fig. 1H). Due to the inability to precisely define bases, the exact nucleotide register remains elusive and, in the modelling, purines or pyrimidines were assigned depending on apparent base size, whilst maintaining Watson-Crick base-pairing in the double-stranded regions. However, since the 40-mer vRNA loop used in the reconstitutions corresponds to 20 nucleotides from the 3′ and 5′ ends of one genome segment, the two observed RNA structures could each correspond to a single vRNA loop with a promoter-like fold, i.e. with a distal duplex (with the loop density missing), and single stranded termini splayed out to bind in the two neighbouring NPs. Less likely, each RNA moiety could result from partial hybridization of two extended vRNAs copies.

Importantly, both the quality of the RNA density in the 2.9 Å resolution focussed map of covering two adjacent NPs from the pseudo-C5 oligomer and the A-from double-stranded RNA helix in the pseudo-C2 oligomer unambiguously reveal the 5′ to 3′ RNA directionality of the RNA bound to TiLV-NP, and this directionality is consistent between all modelled NPs (Supplementary Fig. 8). This key feature is crucial for understanding the NP oligomerization process, not only within the *Amnoonviridae* family but also more broadly within the *Articulavirales* order. Further details will be discussed below.

The RNA binding within TiLV-NP is primarily mediated by (i) interactions between the phosphate backbone and multiple positively charged residues, (ii) base stacking, and (iii) key interactions with the O2′ hydroxyl group, enabling TiLV-NP to selectively bind RNA over DNA (Fig. 4C, D). As mentioned, in the 2.9 Å resolution structure derived from the focussed map of two NPs from the pseudo-C5 oligomer, a total of 12 nucleotides are modelled within one NP (Fig 4C, D). Starting from the 5′ to the 3′ end, the G1 phosphate is stabilised by N225 from the head domain, with its base nestled in a pocket formed by I229, R241 and L285 of the body domain. Weaker density is observed from C2 to A4 as they are not directly stabilised by TiLV-NP but rather stack with each other until reaching U8. A5 and A6 phosphates interact with a triplet of positively charged residues K151, K136 and K139, while A7 ribose O2′ is stabilised by hydrogen bonding with R198 from the head domain. The phosphate group of U8 interacts with K134, and its base is stabilised by N154. U9 takes a sharp turn, repelled by R198, with its base sandwiched between L131, L85 and K83. U10 and A11 phosphates are stabilised by K91, while U10 base is wedged between R198 and F208, which stacks with Y207 to further stabilise A11 base. Finally, U12 phosphate group is surrounded by positively charged residues (R94, K90), while its base stacks with A11 (Fig. 4C, D). Following U12, only disjointed density is observed precluding further RNA model building.

In the pseudo-C2 oligomer structure, continuous density is observed for 37 nucleotides. Within the RNA binding grooves of TiLV-NPi and TiLV-NPi+1, 8 (A2-U9 and A34-G41) nucleotides are stabilised in a similar manner to the pseudo-C5 oligomer (Fig. 4D). However, at the interface of two NPs, the A-form RNA double helix (U10-A33) is formed with the complementary strand extending into the adjacent TiLV-NPi+1 (Fig. 4F, G). In addition to the Watson-Crick base pairing, additional polar residues stabilise the RNA phosphate backbone. Of note, within TiLV-NPi, C20 phosphate is stabilized by R112. In TiLV-NPi+1, R179 and R241 residues stabilise G27 and A30 ribose, with N225, R162, N38 and R158 respectively stabilizing the 5′ phosphates of A30, G31, A32 and A33. A34 stacks on U10 and A33 bases with its phosphate being stabilized by K151, K139 (Fig. 4F, G).

Supporting our structural findings, a previous study used a yeast three-hybrid system to mutate specific positively charged residues of TiLV-NP and assess their impact on RNA binding (22). Mutants “K90A-K91A-R92A-R94A” and K134A significantly disrupted the interaction of TiLV-NP with the RNA. This effect is consistent with our structures, which show that residues K90, K91, R94 and K134 interact with the RNA phosphate backbone, whereas R92 is solvent-exposed and does not interact with the RNA (Fig. 4C). Conversely, substitutions K136A and R158A did not significantly impair RNA binding (22), which agrees with our RNA-bound TiLV-NP structures. K136, though interacting with the phosphate RNA backbone, is surrounded by multiple positively charged residues (K134, K139) that may compensate for its alanine substitution. Similarly, R158, which interacts with the A33 phosphate group in the pseudo-C2 tetramer as part of the A-form RNA double helix, does not interact with RNA in other oligomers. This residue thus likely contributes to the positively charged RNA-binding groove and stabilizes the NP fold, rather than being critical for direct RNA interaction (Fig. 4G).

## DISCUSSION

Numerous apo-NP X-ray structures from orthomyxoviruses, such as influenza A (12,14,15), B (16), D (17), Thogoto (THOV) (18)and ISAV (19)), have been determined as small oligomers (Supplementary Fig. 5). The only X-ray structure with bound RNA is that of FluA/H5N1-NP with 3 nucleotides bound to a basic pocket (15) (Supplementary Fig. 9). Structural studies using cryo-electron microscopy/tomography on purified viral RNPs revealed the overall structures of native FluA (7,20) and THOV (18) RNPs, at nanometer resolution, too low to visualise the RNA. These studies report anti-parallel left-handed assemblies with helical rises of 28 Å and 24.6 Å, and helical twists of -57° to -64° and -55.5°, for Flu and THOV, respectively. More recently, a cryo-EM study at ∼6Å resolution of FluA/H1N1 NP reconstituted with a 12-mer RNA oligomer, revealed a right-handed, super-helical RNP-like structure, but with parallel strands rather than anti-parallel strands expected for a native RNP structures. Low resolution density for 10 to 12 nucleotides was located at the NP-NP interface, but with unspecified directionality (21) (Supplementary Fig. 9). Several functional studies have also identified critical residues in Flu/NP responsible for oligomerization and RNA binding (12) and attempts to model the recruitment of NPs into RNPs have been made (31). However, no prior published studies have provided insights into how RNA binds within the positively charged groove of NP, nor into the RNA 5′ to 3′ directionality, which is crucial for understanding NP oligomerization during the articulaviral replication process.

Here we present the biophysical and structural characterization of RNA-bound oligomers of the prototypic amnoonviral NP from TiLV. We show that, in vitro, apo TiLV-NP primarily forms tetramers and trimers, and in presence of RNA, oligomerizes into higher-order closed, planar oligomers, including pentamers, hexamers and larger assemblies, with each NP bound to RNA (Fig. 1). The TiLV-NP sequence is only similar to putative NPs from other Amnoonviruses. The most closely related (90% homology) is the recently reported Fancy Tailed Guppy fish virus segment 4 protein (32) (Supplementary Fig. 10). Much more distant sequences are from Flavolineata, Namensis and Asotus viruses (33), which show the most conservation between residues 230 and 305 (TiLV-NP numbering), comprising part of the body domain, the hinges and oligomerisation loop (Supplementary Fig. 10). Despite the complete divergence of sequence and having a significantly reduced size compared to classical orthomyxovirus NPs, TiLV-NP maintains the same fold and architecture. It comprises body and head domains, along with a protruding oligomerization loop flanked by flexible hinges (Fig. 2). These conserved structural elements form a positively charged RNA binding groove and a hydrophobic pocket that mediates NP oligomerization (Fig. 3). In addition, we describe for the first time an orthomyxovirus-like NP structure bound to RNA and reveal the 5′ to 3′ RNA directionality (Fig. 4). This is unambiguously supported by both the high-resolution of the cryo-EM maps and the presence of an A-form RNA double helix at the NP-NP interface of the pseudo-C2 oligomer. These results provide the first insight into how the RNA is encapsidated within circular oligomers, starting at the 5′ end and progressively covering the entire RNA. The presence of an A-form RNA double helix at the NP-NP interface also highlights that inter-NP flexibility allows accommodation of RNA secondary structures. Secondary structures have been detected in influenza A vRNA (34-36), as well as the irregular positioning of NP along the RNP (37). These phenomena have been postulated to mediate intersegment RNA-RNA interactions and be implicated in selective bundling of influenza vRNPs prior to packaging in budding virions Furthermore, superposition of the RNA-bound TiLV-NP structure onto FluA/H1N1- and H5N1-NP suggests that the general RNA-binding mode is likely conserved within articulaviral NPs (Supplementary Fig. 9). The superposed RNA is well integrated within the FluA/NP RNA binding groove, with previously described RNA-interacting residues clustering nearby (Supplementary Fig. 9). We also note that the RNA directionality we observe is consistent with the structure of FluA/H5N1-NP with 3 nucleotides bound (15) (Supplementary Fig. 9). It is also consistent with the ‘tail-loop first’ model of NP directionality deduced using oligomerisation defective mutants in cells (31) (Supplementary Fig. 9, 11).

Nevertheless, several questions remain. Although previous studies have provided low-resolution EM images of purified TiLV RNPs (22), their quality precluded further characterization, and the helical parameters of TiLV-NC remain unknown. Assuming TiLV-NC adopts an antiparallel superhelix like influenza, it is tempting to suggest that the circularized TiLV-NP pentamers and hexamers may correspond to the loop found at one extremity of the RNP or could, perhaps, correspond to a nascent RNP formed within the replication complex once the RNA product is beginning to bulging out (10,11). In the pseudo-C2 structure, the A-form RNA that splays out into single-stranded 3’ and 5’ extensions that bind to adjacent NPs, could also mimic the polymerase proximal promoter duplex separating into incoming and outgoing strands, each binding an NP at the start of the RNP superhelix. Indeed, superposition of the TiLV-NP A-from RNA onto the distal duplex seen in TiLV polymerase (4) shows that both NPs avoid severe steric clashes with TiLV-Pol (Supplementary Fig. 11). However, in vitro reconstitution of TiLV RNPs, as recently achieved for Flu (21), or cryo-ET on purified viral RNPs, will be necessary to provide further insights into TiLV RNP morphology and dynamics during RNA synthesis.

Finally, despite having mapped the previously described mutations introduced using a yeast-three hybrid system onto our TiLV-NP structures and showing consistency of results, in *cellulo* functional studies remain challenging due to the lack of an established TiLV minigenome system. Very recently, a reverse genetics system for TiLV rescue has been described (38), the first step to more flexible laboratory tools to probe the mechanism of TiLV replication and establishing a platform for investigating antiviral strategies targeting either the NP-NP, NP-RNA interfaces or the polymerase function.

To conclude, studying phylogenetically divergent viruses such as TiLV offers valuable insights into unravelling conserved or divergent mechanisms of transcription and replication within the *Articulavirales* order.

## DATA AVAILABILITY

The data that support this study are available from the corresponding authors upon reasonable request. The EM maps and co-ordinates generated in this study have been deposited in the Electron Microscopy Data Bank and the Protein Data Bank (summarised in Supplementary Tables 1-4):

TiLV-NP pentamer (pseudo-C5) (local refinement around 2 TiLV-NPs) PDB ID 9HBR EMD-52027.

TiLV-NP pentamer (pseudo-C5) PDB ID 9HBT EMD-52029.

TiLV-NP tetramer (pseudo-C2) (local refinement around 2 TiLV-NPs) PDB ID 9HBU EMD-52030.

TiLV-NP tetramer (pseudo-C2) PDB ID 9HBS EMD-52028.

TiLV-NP tetramer (pseudo-C4) (local refinement around 2 TiLV-NPs) PDB ID 9HBV EMD-52031.

TiLV-NP tetramer (pseudo-C4) PDB ID 9HBW EMD-52032.

TiLV-NP hexamer (pseudo-C6) (local refinement around 2 TiLV-NPs) PDB ID 9HBX EMD-52033.

TiLV-NP hexamer (pseudo-C6) (local refinement around 3 TiLV-NPs) PDB ID 9HBY EMD-52035.

TiLV-NP hexamer (pseudo-C6) PDB ID 9HBZ EMD-52035.

Source data are provided in a Supplemental file associated with this paper.

## SUPPLEMENTARY DATA

Supplementary Data are available at NAR online.

## AUTHOR CONTRIBUTIONS

SC and BA conceived the project. BA, MP and KH performed cloning, protein expression and purification. BA performed all in vitro biochemical, biophysical and cryo-EM analyses. SC and BA did model building and refinement. BA and SC prepared the manuscript.

## ACKNOWLEDGEMENTS

We thank Romain Linares and Iskander Khusainov for access to the EMBL Grenoble Glacios; Romain Linares and Joseph Bartho for data collections performed on the Titan Krios at EMBL Heidelberg; Aymeric Peuch for support using the joint EMBL-IBS computer cluster; Caroline Mas for assistance and access to the biophysical platform. This work used the platforms of the Grenoble Instruct-ERIC center (ISBG; UAR 3518 CNRS-CEA-UGA-EMBL) within the Grenoble Partnership for Structural Biology (PSB), supported by FRISBI (ANR-10-INBS-0005-02) and GRAL, financed within the University Grenoble Alpes graduate school (Ecoles Universitaires de Recherche) CBH-EUR-GS (ANR-17-EURE-0003).

## Supplementary Data

**Supplementary Table 1.** Cryo-EM data collection, refinement and validation statistics for the RNA-bound TiLV-NP pseudo-C5 oligomer structures.

| Sample | TiLV-NP with 40-mer vRNA<br>2 NPs from pentamer | TiLV-NP with 40-mer vRNA<br>Complete pentamer |
| --- | --- | --- |
| PDB ID | 9HBR | 9HBT |
| EMDB ID | EMD-52027 | EMD-52029 |
| Data collection and processing |  |  |
| Microscope | ThermoFisher TEM Titan Krios G3 |  |
| Voltage (kV) | 300 |  |
| Camera | Gatan K3 / BioQuantum EF |  |
| Magnification | 105,000 |  |
| Nominal defocus range (μm) | -0.8 / -2.0 |  |
| Electron exposure (e- /Å²) | 40 |  |
| Pixel size (Å) | 0.822 |  |
| Initial micrographs (no.) | 2330 |  |
| Final micrographs (no.) | 2067 |  |
| Refinement |  |  |
| Particles per class (no.) | 225,425 | 352,053 |
| Map resolution (Å),<br>0.143 FSC | 2.90 | 3.46 |
| Model resolution (Å),<br>0.5 FSC | 3.0 | 3.7 |
| Map sharpening *B* factor (Å²) | -116 | -127 |
| Map versus model cross-correlation (CCmask) | 0.8105 | 0.6841<br>(2 subunits poor density) |
| Model composition |  |  |
| Non-hydrogen atoms | 5268 | 13417 |
| Protein residues | 625 | 1581 |
| RNA nts | 22 | 57 |
| Mean *B* factors (Å²) |  |  |
| Protein | 69.63 | 58.98 |
| RNA | 125.79 | 97.53 |
| RMS deviations |  |  |
| Bond lengths (Å) | 0.003 | 0.002 |
| Bond angles (°) | 0.397 | 0.382 |
| Validation |  |  |
| MolProbity score | 1.23 | 1.50 |
| All-atom clash score | 1.54 | 3.67 |
| Poor rotamers (%) | 2.76 | 2.81 |
| Ramachandran plot |  |  |
| Favored (%) | 98.21 | 98.28 |
| Allowed (%) | 1.79 | 1.72 |
| Outliers (%) | 0.0 | 0.0 |

**Supplementary Table 2.**
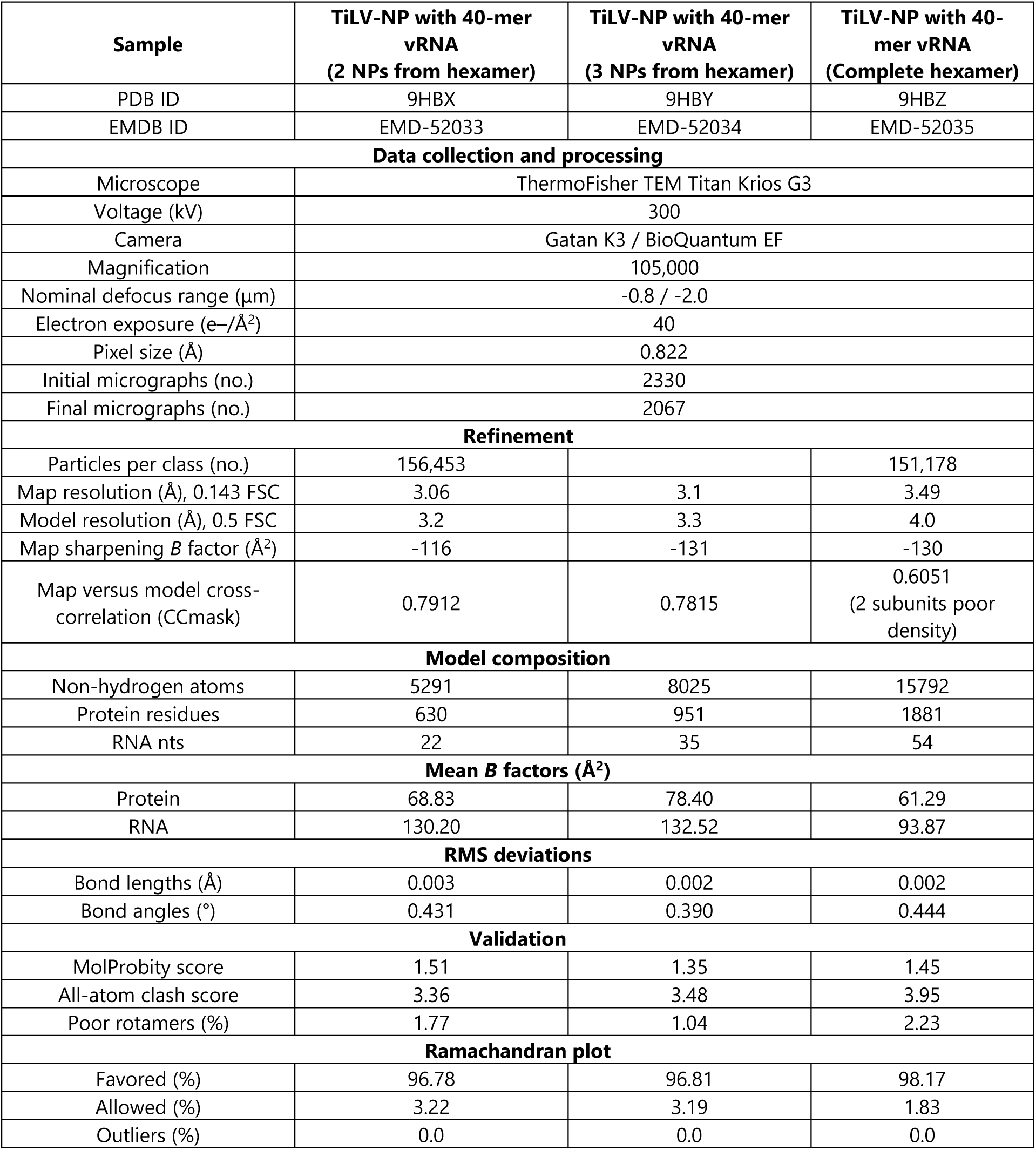
Cryo-EM data collection, refinement and validation statistics for the RNA-bound TiLV-NP pseudo-C6 oligomer structures.

**Supplementary Table 3.**
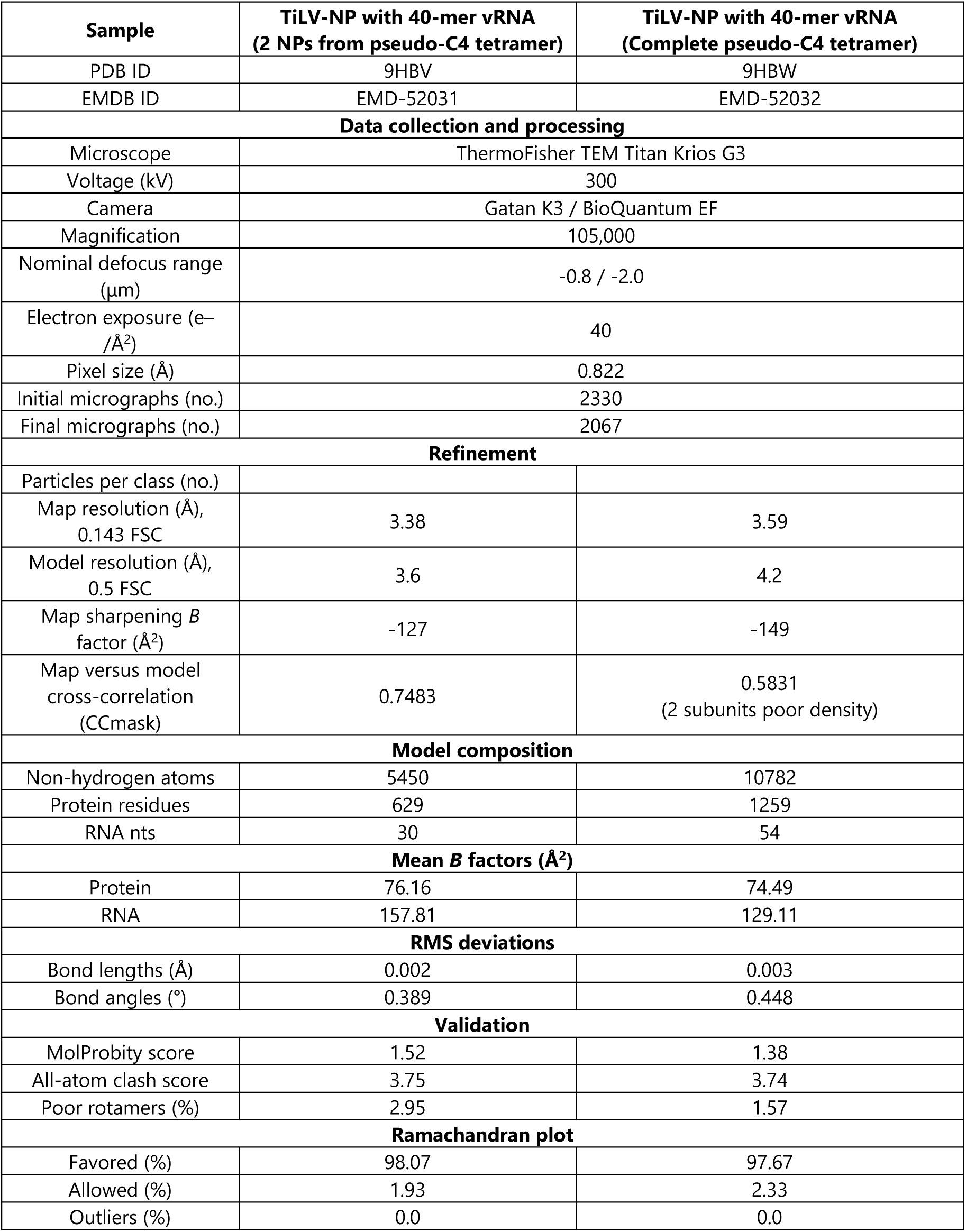
Cryo-EM data collection, refinement and validation statistics for the RNA-bound TiLV-NP pseudo-C4 oligomer structures.

| Sample | TiLV-NP with 40-mer vRNA<br>(2 NPs from pseudo-C4 tetramer) | TiLV-NP with 40-mer vRNA<br>(Complete pseudo-C4 tetramer) |
| --- | --- | --- |
| PDB ID | 9HBV | 9HBW |
| EMDB ID | EMD-52031 | EMD-52032 |
| Data collection and processing |  |  |
| Microscope | ThermoFisher TEM Titan Krios G3 |  |
| Voltage (kV) | 300 |  |
| Camera | Gatan K3 / BioQuantum EF |  |
| Magnification | 105,000 |  |
| Nominal defocus range (μm) | -0.8 / -2.0 |  |
| Electron exposure (e-/Å²) | 40 |  |
| Pixel size (Å) | 0.822 |  |
| Initial micrographs (no.) | 2330 |  |
| Final micrographs (no.) | 2067 |  |
| Refinement |  |  |
| Particles per class (no.) |  |  |
| Map resolution (Å), 0.143 FSC | 3.38 | 3.59 |
| Model resolution (Å), 0.5 FSC | 3.6 | 4.2 |
| Map sharpening *B* factor (Å²) | -127 | -149 |
| Map versus model cross-correlation (CCmask) | 0.7483 | 0.5831<br>(2 subunits poor density) |
| Model composition |  |  |
| Non-hydrogen atoms | 5450 | 10782 |
| Protein residues | 629 | 1259 |
| RNA nts | 30 | 54 |
| Mean *B* factors (Å²) |  |  |
| Protein | 76.16 | 74.49 |
| RNA | 157.81 | 129.11 |
| RMS deviations |  |  |
| Bond lengths (Å) | 0.002 | 0.003 |
| Bond angles (°) | 0.389 | 0.448 |
| Validation |  |  |
| MolProbity score | 1.52 | 1.38 |
| All-atom clash score | 3.75 | 3.74 |
| Poor rotamers (%) | 2.95 | 1.57 |
| Ramachandran plot |  |  |
| Favored (%) | 98.07 | 97.67 |
| Allowed (%) | 1.93 | 2.33 |
| Outliers (%) | 0.0 | 0.0 |

**Supplementary Table 4.**
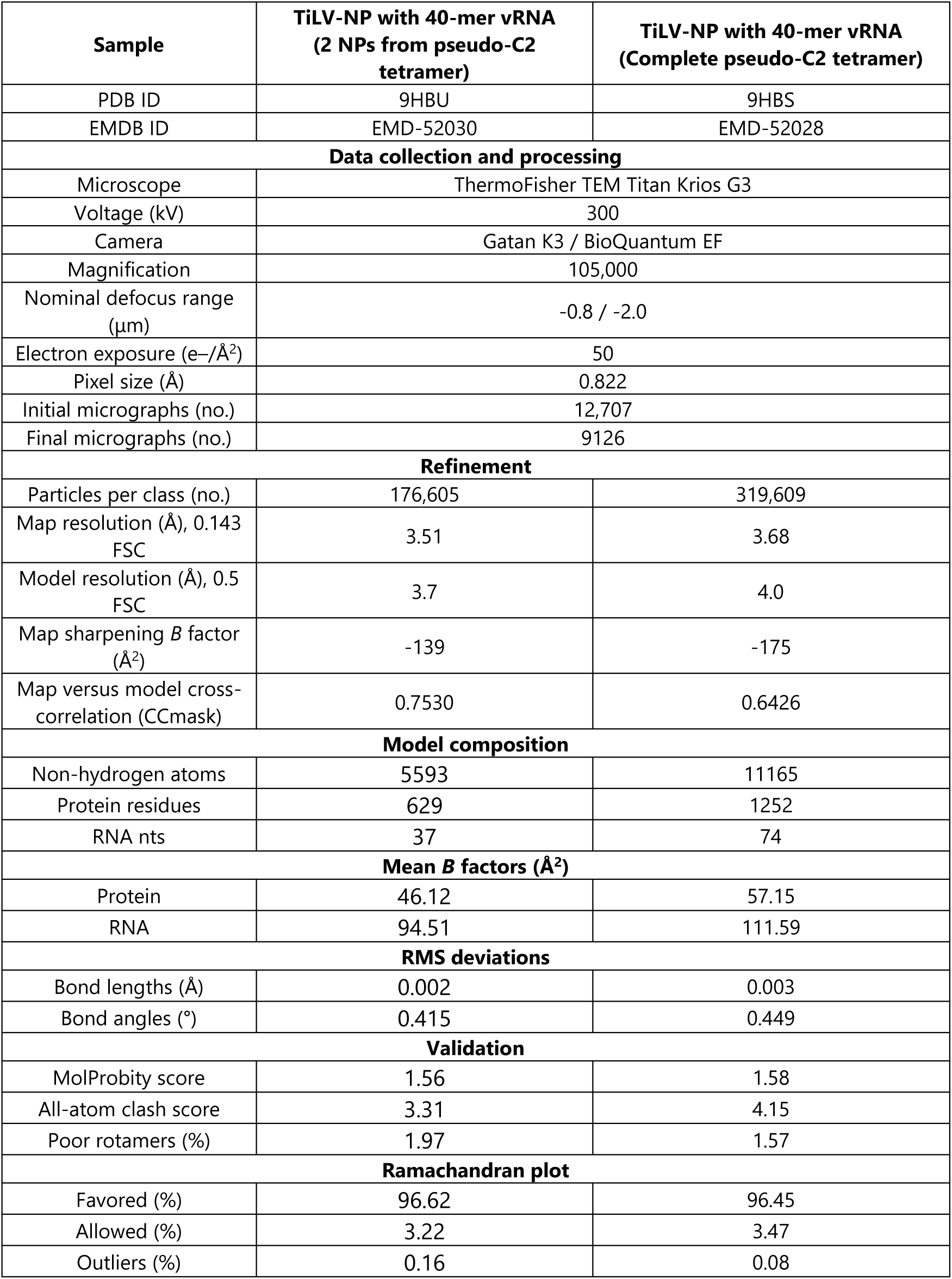
Cryo-EM data collection, refinement and validation statistics for the RNA-bound TiLV-NP pseudo-C2 oligomer structures.

| Sample | TiLV-NP with 40-mer vRNA<br>(2 NPs from pseudo-C2 tetramer) | TiLV-NP with 40-mer vRNA<br>(Complete pseudo-C2 tetramer) |
| --- | --- | --- |
| PDB ID | 9HBU | 9HBS |
| EMDB ID | EMD-52030 | EMD-52028 |
| Data collection and processing |  |  |
| Microscope | ThermoFisher TEM Titan Krios G3 |  |
| Voltage (kV) | 300 |  |
| Camera | Gatan K3 / BioQuantum EF |  |
| Magnification | 105,000 |  |
| Nominal defocus range (μm) | -0.8 / -2.0 |  |
| Electron exposure (e-/Å²) | 50 |  |
| Pixel size (Å) | 0.822 |  |
| Initial micrographs (no.) | 12,707 |  |
| Final micrographs (no.) | 9126 |  |
| Refinement |  |  |
| Particles per class (no.) | 176,605 | 319,609 |
| Map resolution (Å), 0.143 FSC | 3.51 | 3.68 |
| Model resolution (Å), 0.5 FSC | 3.7 | 4.0 |
| Map sharpening *B* factor (Å²) | -139 | -175 |
| Map versus model cross-correlation (CCmask) | 0.7530 | 0.6426 |
| Model composition |  |  |
| Non-hydrogen atoms | 5593 | 11165 |
| Protein residues | 629 | 1252 |
| RNA nts | 37 | 74 |
| Mean *B* factors (Å²) |  |  |
| Protein | 46.12 | 57.15 |
| RNA | 94.51 | 111.59 |
| RMS deviations |  |  |
| Bond lengths (Å) | 0.002 | 0.003 |
| Bond angles (°) | 0.415 | 0.449 |
| Validation |  |  |
| MolProbity score | 1.56 | 1.58 |
| All-atom clash score | 3.31 | 4.15 |
| Poor rotamers (%) | 1.97 | 1.57 |
| Ramachandran plot |  |  |
| Favored (%) | 96.62 | 96.45 |
| Allowed (%) | 3.22 | 3.47 |
| Outliers (%) | 0.16 | 0.08 |

**Supplementary Figure 1.**
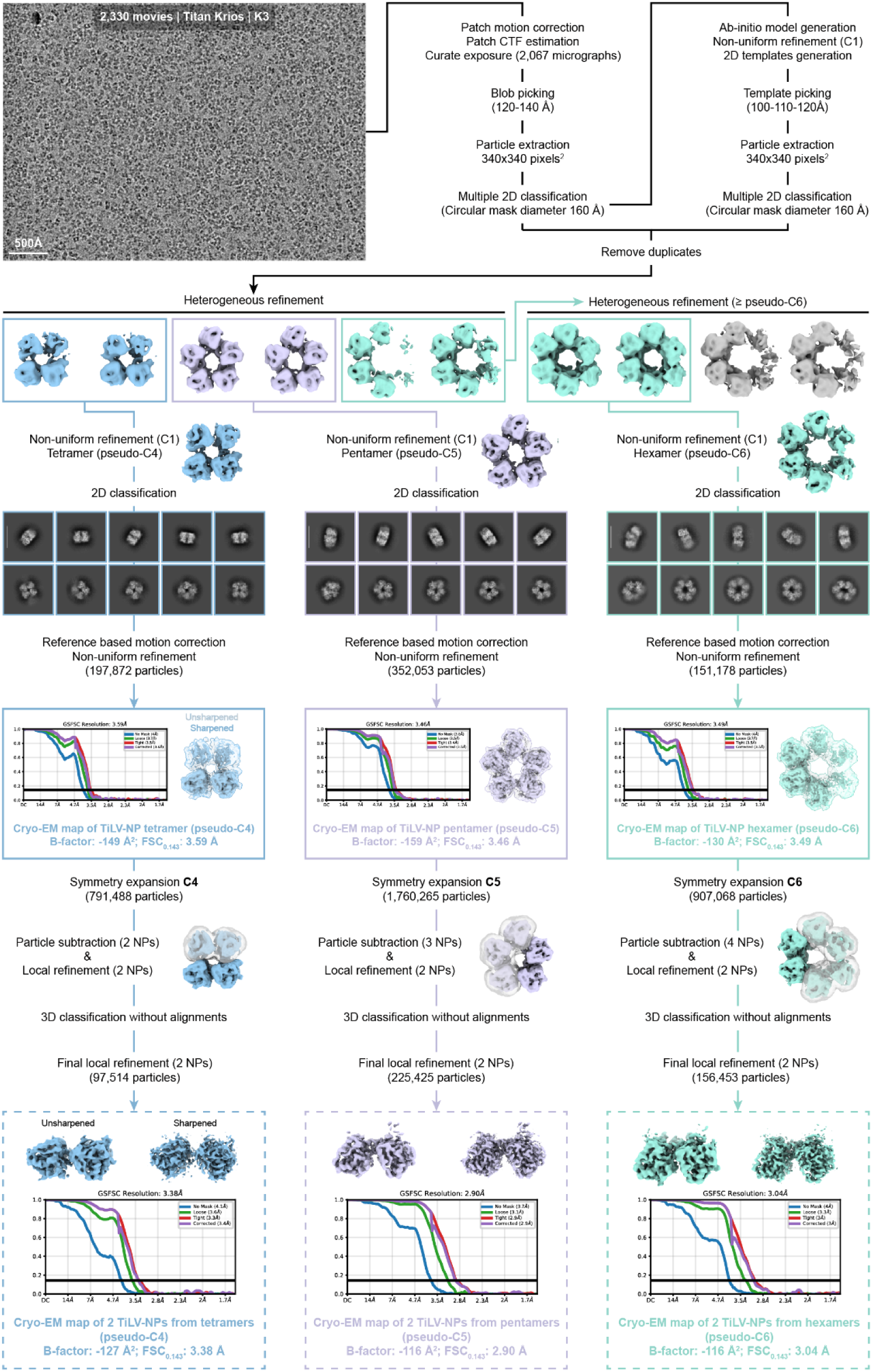
General cryo-EM image processing strategy applied to obtain TiLV-NP structures (tetrameric pseudo-C4, pentameric pseudo-C5, and hexameric pseudo-C6). Schematics of the image processing strategy used with the data collected on a Titan Krios equipped with a Gatan K3 direct electron detector mounted on a Gatan Bioquantum energy filter. Representative realigned micrograph (-3 µM defocus, low-pass filtered at 5 Å, scale bar = 500 Å), 2D class averages (scale bar = 120 Å), and 3D maps are displayed. Fourier shell correlation curves (FSC) are shown, and the overall resolution based on the FSC0.143 criteria indicated.

**Supplementary Figure 2.**
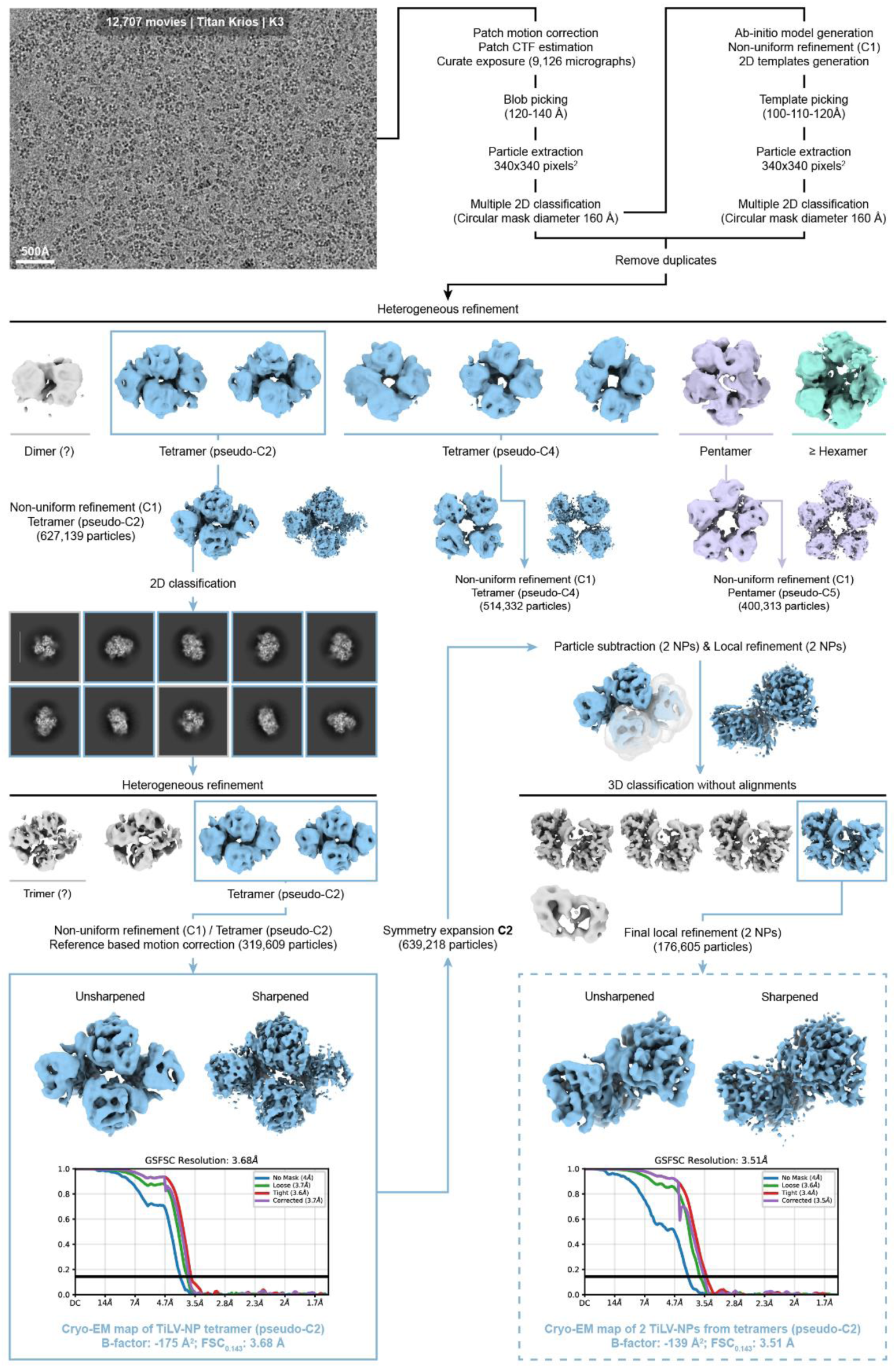
General cryo-EM image processing strategy applied to obtain TiLV-NP structures (tetrameric pseudo-C2). Schematics of the image processing strategy used with the data collected on a Titan Krios equipped with a Gatan K3 direct electron detector mounted on a Gatan Bioquantum energy filter. Representative realigned micrograph (-3 µM defocus, low-pass filtered at 5 Å, scale bar = 500 Å), 2D class averages (scale bar = 120 Å), and 3D maps are displayed. Fourier shell correlation curves (FSC) are shown, and the overall resolution based on the FSC0.143 criteria indicated.

**Supplementary Figure 3.**
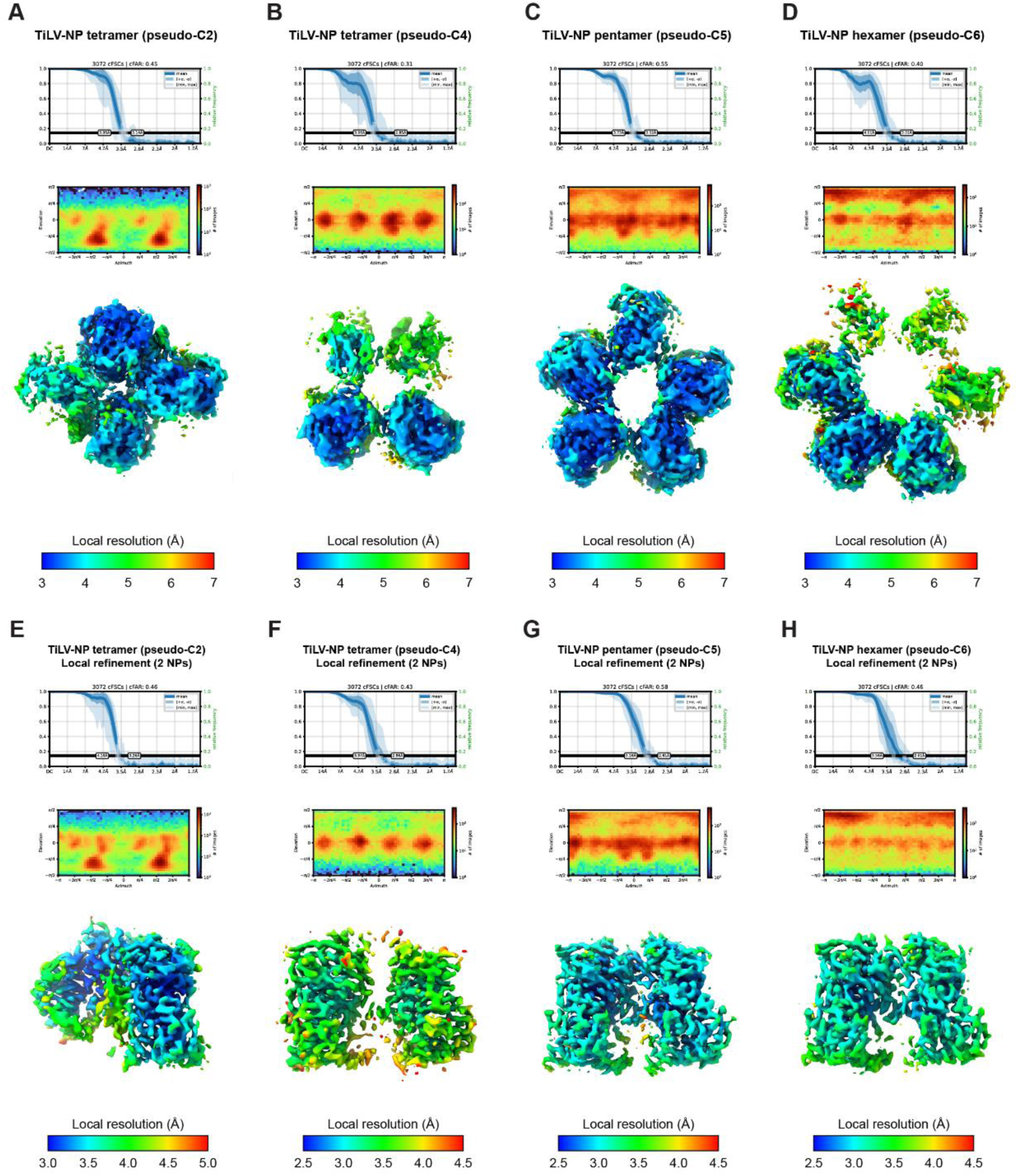
3D-FSCs, orientation distribution and local resolution of each TiLV-NP structures. A-D. 3D-FSCs, orientation distribution and local resolution of each complete TiLV-NP structures (A) pseudo-C2; (B) pseudo-C4; (C) pseudo-C5; (D) pseudo-C6. E-H. 3D-FSCs, orientation distribution and local resolution of each local refinement around two TiLV-NPs for each oligomer (E) pseudo-C2; (F) pseudo-C4; (G) pseudo-C5; (H) pseudo-C6.

**Supplementary Figure 4.**
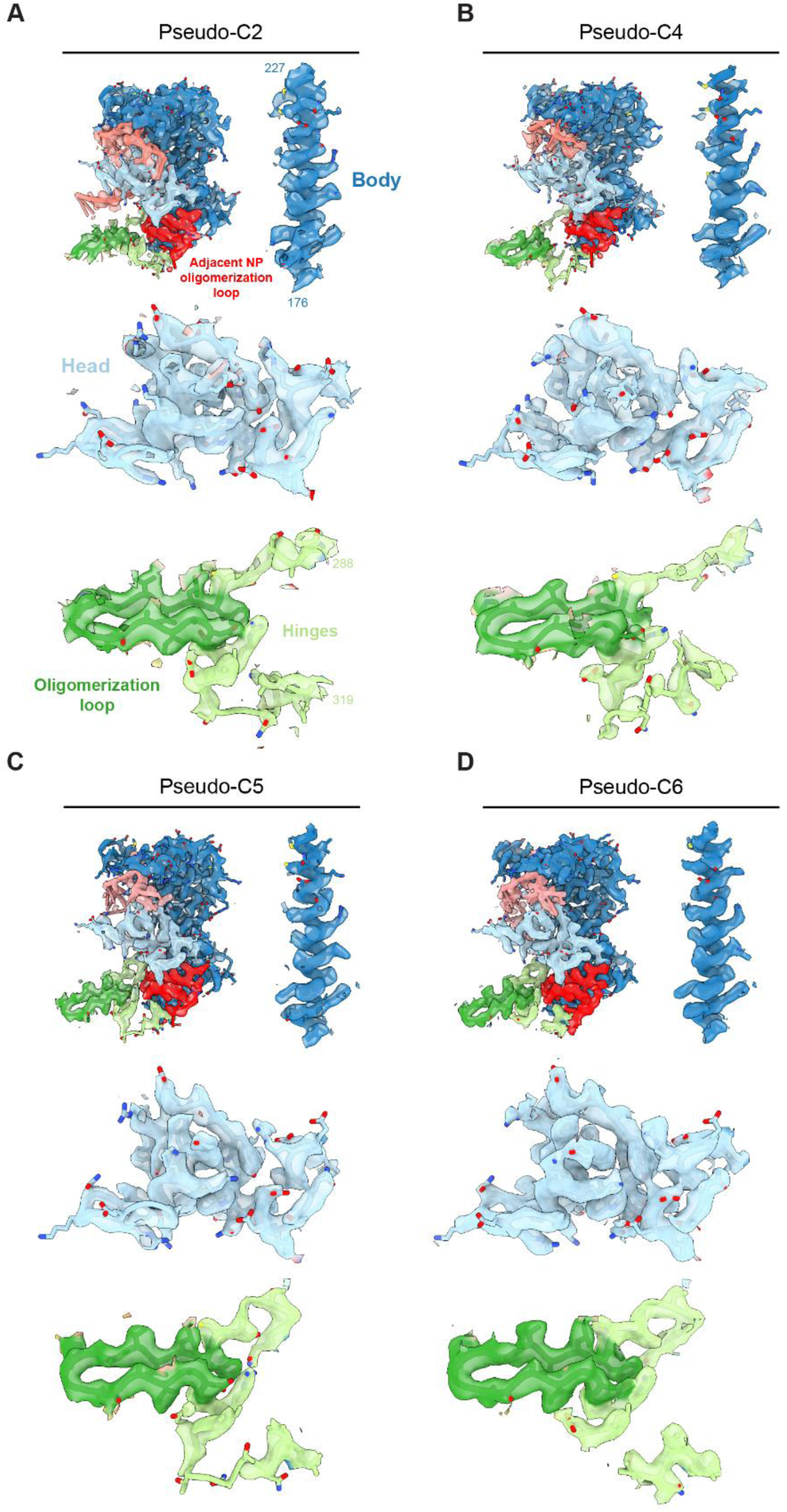
Cryo-EM map quality of TiLV-NP structures. A-D. Extracted coulomb potential of a single TiLV-NP from the local refinement around two NPs from the (A) pseudo-C2 tetramer, (B) pseudo-C4 tetramer, (C) pseudo-C5 pentamer, (D) pseudo-C6 hexamer maps. Additional EM densities are shown for the body, head, hinges, and the oligomerization loop, with side chains clearly visible.

**Supplementary Figure 5.**
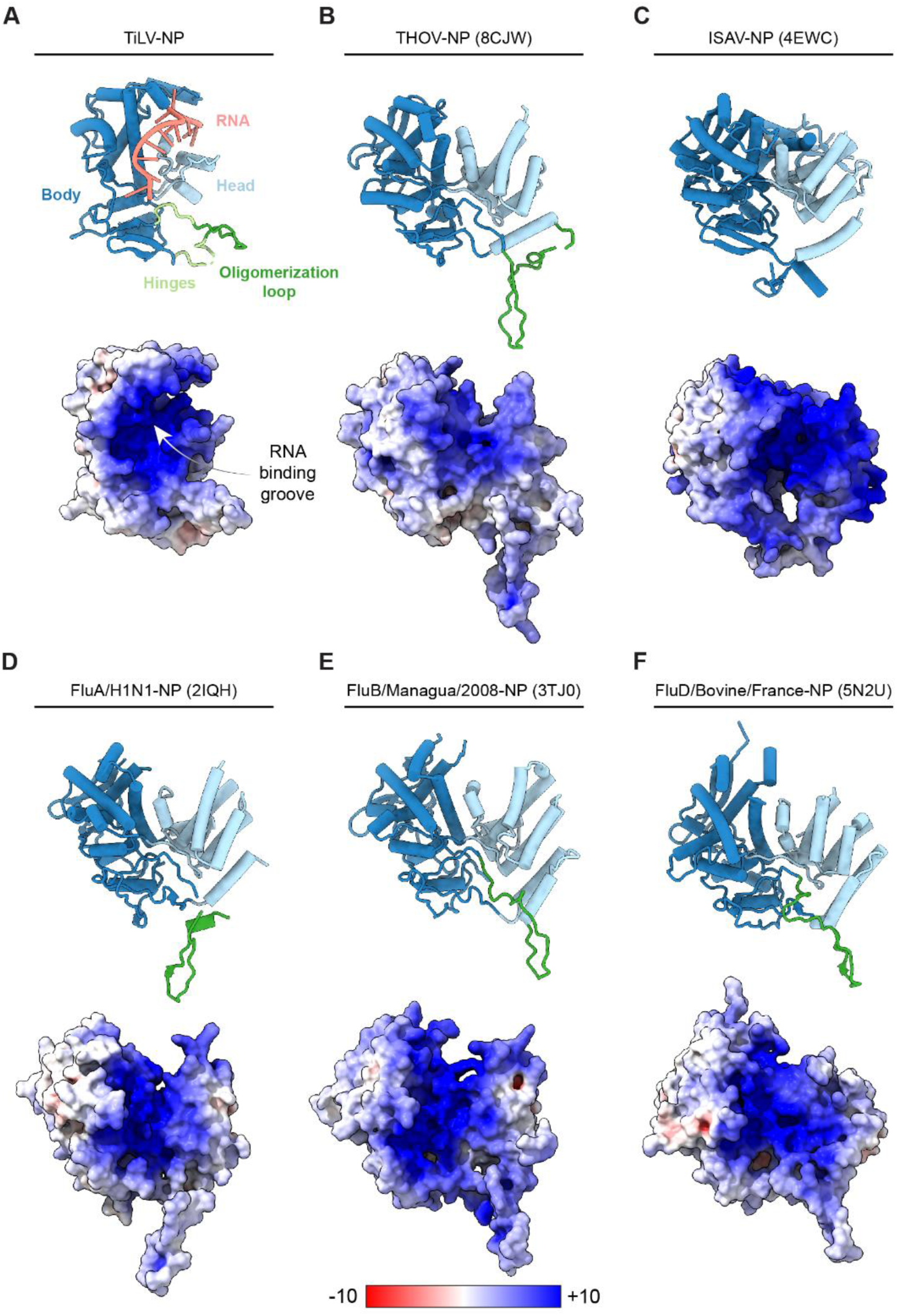
Structural comparison of TiLV-NP monomer with other orthomyxo(-like) viruses NPs. A. Top: Cartoon representation of RNA-bound TiLV-NP. TiLV-NP domains and RNA are coloured as in Fig. 2. Bottom: Surface representation of RNA-bound TiLV-NP coloured based on electrostatic potential. B. Top: Cartoon representation of THOV-NP (PDB 8CJW). THOV-NP domains are coloured as TiLV-NP. Bottom: Surface representation of THOV-NP coloured based on electrostatic potential. C. Top: Cartoon representation of ISAV-NP (PDB 4EWC). ISAV-NP domains are coloured as TiLV-NP. Bottom: Surface representation of ISAV-NP coloured based on electrostatic potential. D. Top: Cartoon representation of FluA/H1N1-NP (PDB 2IQH). FluA/H1N1-NP domains are coloured as TiLV-NP. Bottom: Surface representation of FluA/H1N1-NP coloured based on electrostatic potential. E. Top: Cartoon representation of FluB/Managua/2008-NP (PDB 3TJ0). FluB/Managua/2008-NP domains are coloured as TiLV-NP. Bottom: Surface representation of FluB/Managua/2008-NP coloured based on electrostatic potential. F. Top: Cartoon representation of FluD/Bovine/France-NP (PDB 5N2U). FluD/Bovine/France-NP domains are coloured as TiLV-NP. Bottom: Surface representation of FluD/Bovine/France-NP coloured based on electrostatic potential.

**Supplementary Figure 6.**
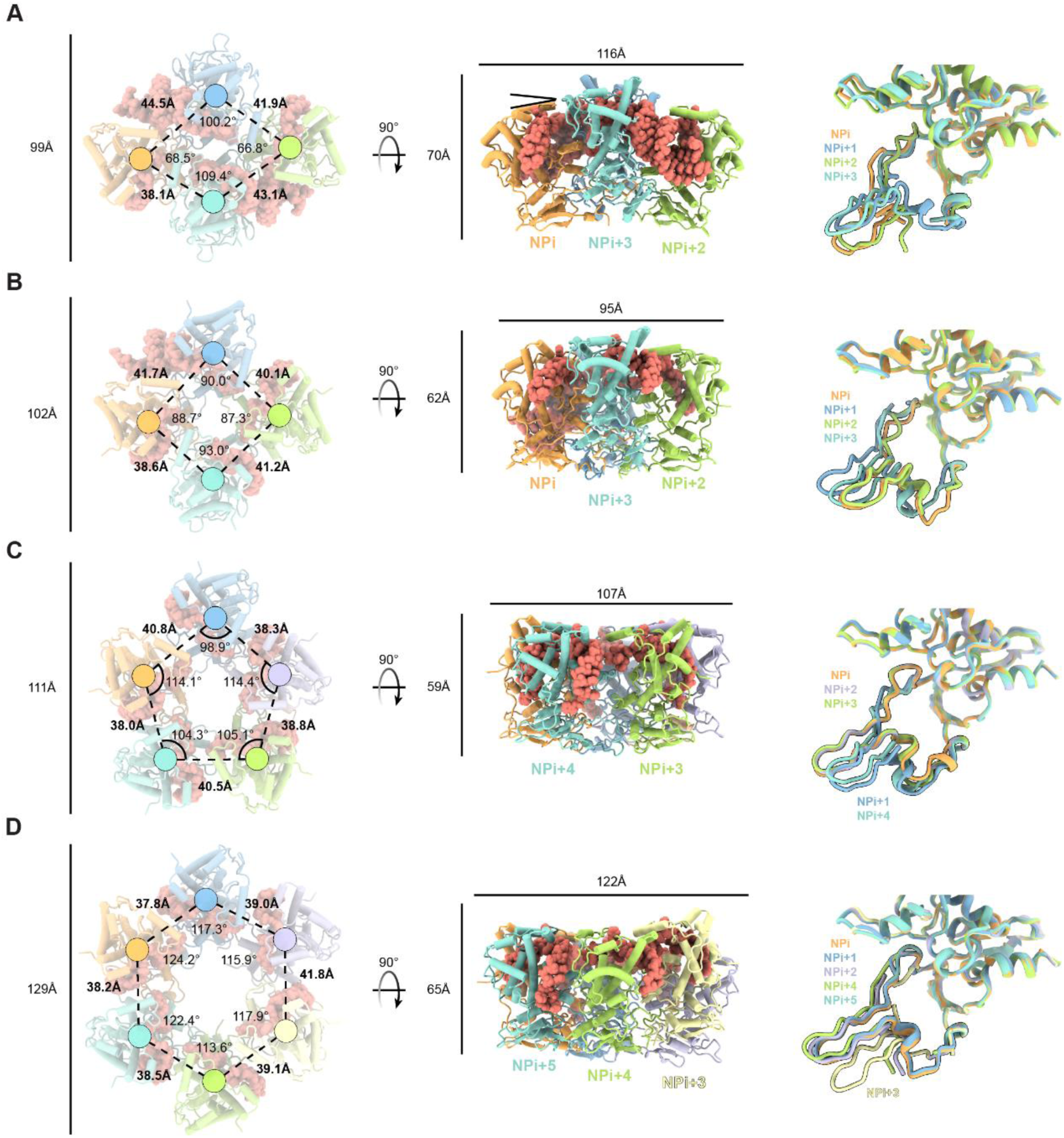
Pseudo-symmetrical arrangement of TiLV-NP oligomers. A-D. Cartoon representation of (A) TiLV-NP tetramer (pseudo-C2), (B) TiLV-NP tetramer (pseudo-C4), (C) TiLV-NP pentamer (pseudo-C5), (D) TiLV-NP hexamer (pseudo-C6), with NP centroids shown as spheres. Oligomer dimensions, inter-NP distances and angles are indicated. Superposition of individual NPs for each oligomer highlights flexibility in the hinges and the oligomerization loop, underlined with a black line.

**Supplementary Figure 7.**
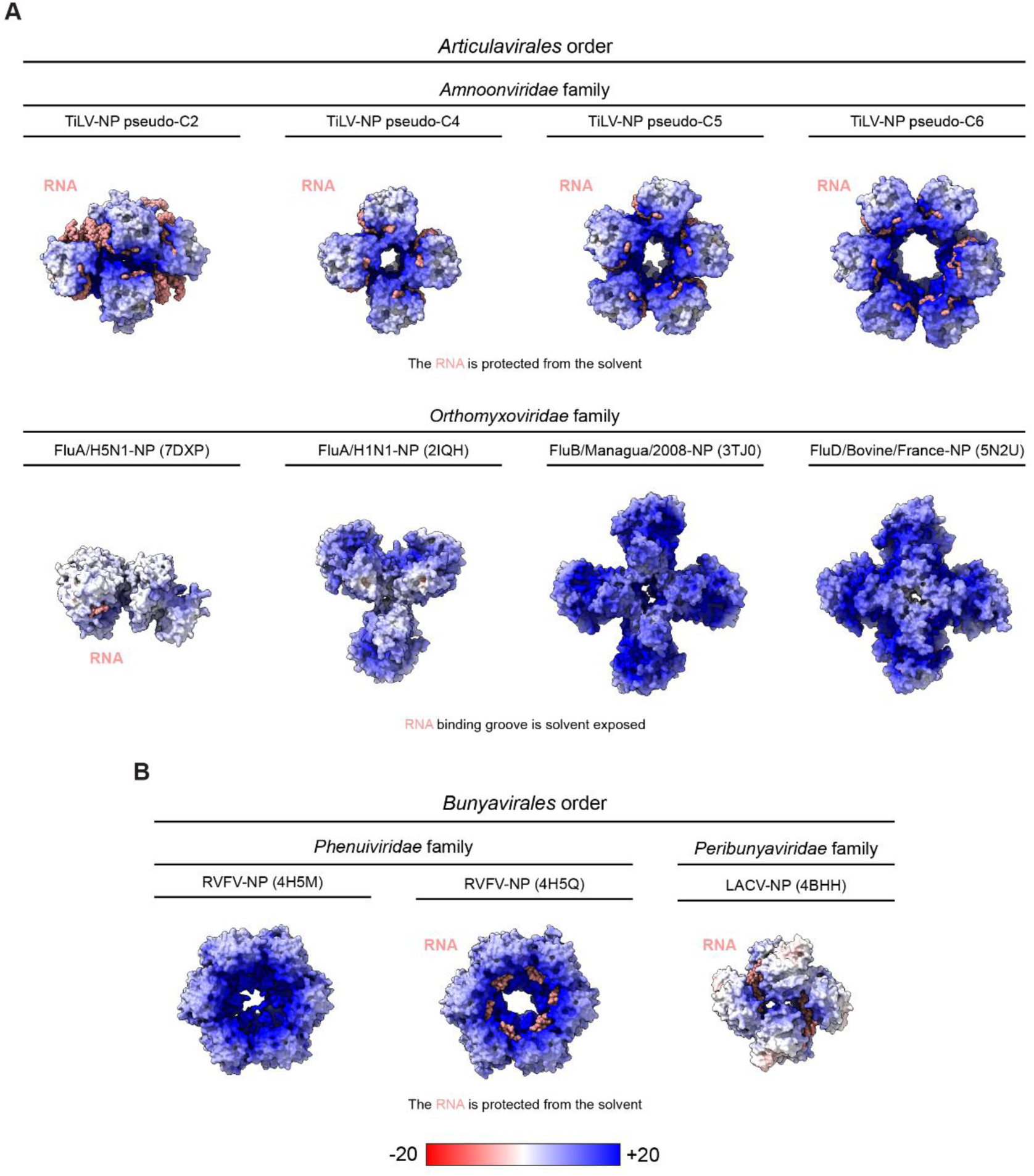
Structural comparison of TiLV-NP oligomers with orthomyxo- and bunya-virus NPs. A. Surface representations of NPs from orthomyxo(-like) viruses, coloured based on electrostatic potential. The top row shows RNA-bound TiLV-NP structures determined by cryo-EM in this study. The bottom row shows X-ray structures of FluA/H5N1-NP (PDB 7DXP), FluA/H1N1-NP (PDB 2IQH), FluB/Managua/2008-NP (PDB 3TJ0), FluD/Bovine/France-NP (PDB 5N2U). In TiLV-NP oligomers, the RNA is protected from the solvent, whereas in the Flu-NP oligomer X-ray structures, the RNA binding groove is solvent exposed. The RNA is displayed as spheres, coloured in salmon. B. Surface representations of bunyavirus NPs X-ray structures (Rift Valley Fever virus “RVFV”, PDB 4H5M / 4H5Q; La Crosse virus “LACV”, PDB 4BHH), coloured based on electrostatic potential. The RNA binding groove is protected from the solvent, similar to TiLV oligomers.

**Supplementary Figure 8.**
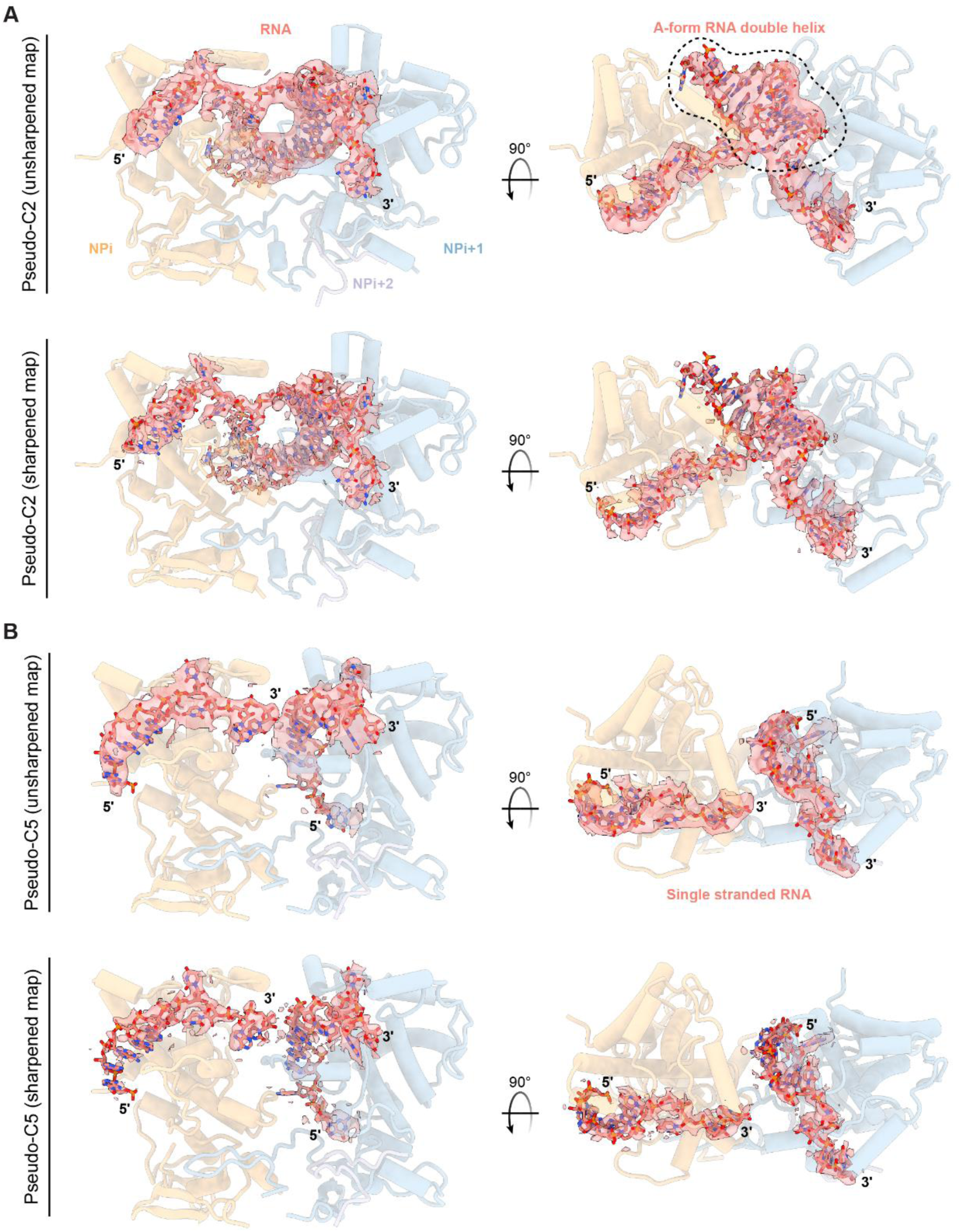
Cryo-EM map quality of the RNA within TiLV-NP structures. A. RNA coulomb potential extracted from the local refinement around two TiLV NPs from the pseudo-C2 tetramer map. The RNA is coloured in salmon, NPi orange, NPi+1 blue and NPi+2 mauve. The 5′ and 3′ ends are annotated. Both unsharpened and sharpened maps are shown. B. RNA coulomb potential extracted from the local refinement around two TiLV NPs from the pseudo-C5 pentamer map. The RNA is coloured in salmon, NPi orange, NPi+1 blue and NPi+2 mauve. The 5′ and 3′ ends are annotated. Both unsharpened and sharpened maps are shown.

**Supplementary Figure 9.**
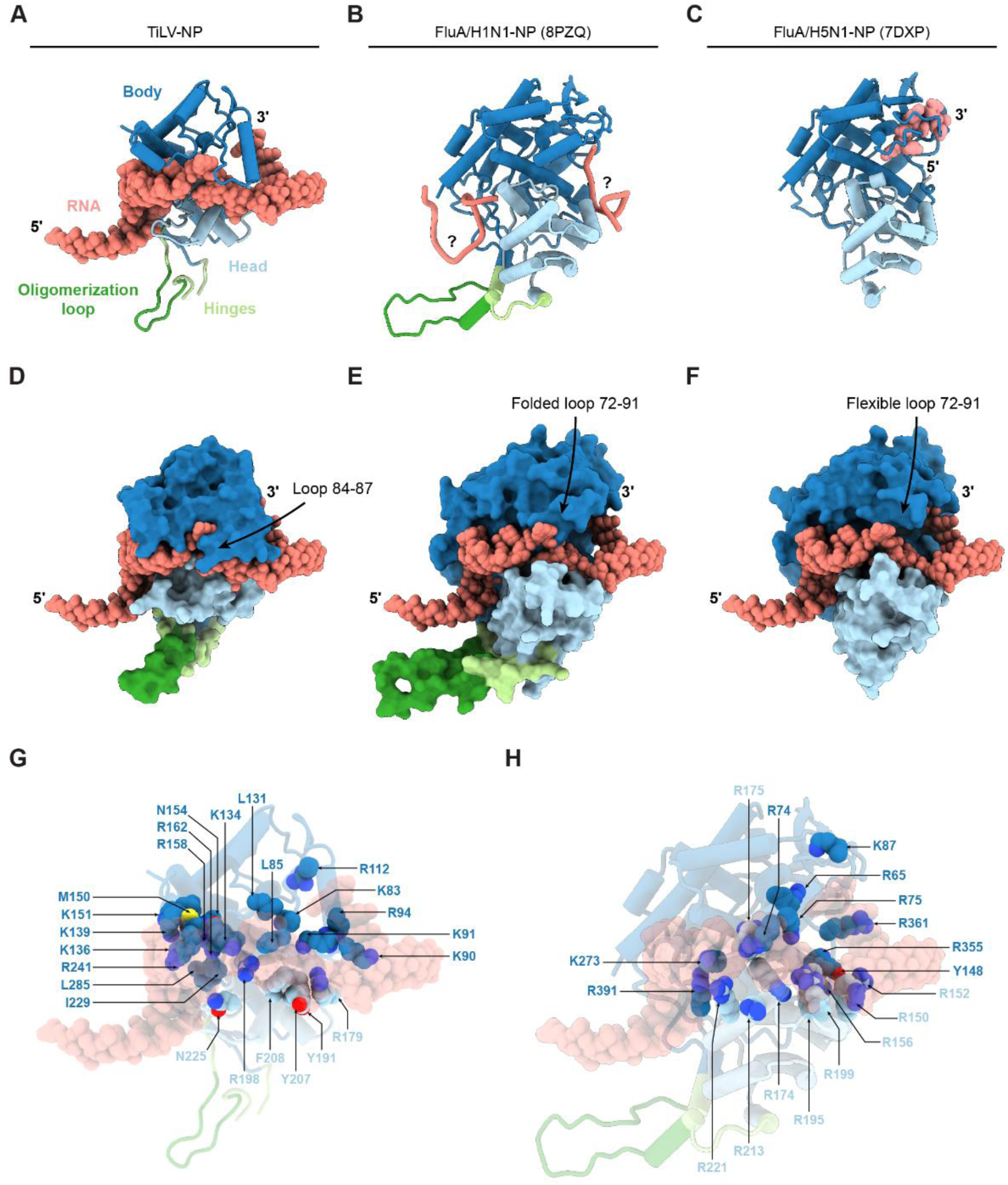
Superposition of TiLV-NP RNA on influenza NPs. A. Cartoon representation of RNA-bound TiLV-NP. All RNA conformations observed in the TiLV-NP pseudo-C2 and pseudo-C5 structures are superimposed (see Fig. 4B, F). The RNAs are shown as spheres. TiLV-NP domains are coloured as in Fig 2. The 5′ and 3′ ends are annotated. B. Cartoon representation of FluA/H5N1-NP (PDB 8PZQ). Only one NP is displayed (chain A). The RNA phosphate backbone is shown as liquorice as the RNA polarity and base position are unknown. Flu-NP domains are coloured as in Fig 2. C. Cartoon representation of FluA/H1N1-NP (PDB 7DXP). Only one NP is displayed (chain B). The RNA with methylated 2′ OHs is shown as spheres. The 5′ and 3′ ends are annotated. Flu-NP domains are coloured as in Fig 2. 1. D. Similar to A, but TiLV-NP is displayed as surface. 2. E. Superposition of TiLV-NP RNAs onto FluA/H5N1-NP (PDB 8PZQ). TiLV-NP RNAs follow the Flu-NP RNA binding groove. One “clash” is next to the Flu-NP folded loop 72-91, which is flexible in FluA/H1N1-NP (PDB 7DXP). 1. F. Superposition of TiLV-NP RNAs onto FluA/H1N1-NP (PDB 7DXP). TiLV-NP RNAs follow the Flu-NP RNA binding groove. The RNA polarity is conserved with the 3 methylated nucleotides observed in FluA/H1N1-NP (PDB 7DXP) X-ray structure. 2. G. Summary of TiLV-NP residues involved in RNA interactions. TiLV-NP and all superimposed RNA conformations observed in the TiLV-NP pseudo-C2 and pseudo-C5 structures are transparent. Key interacting residues are displayed as spheres and coloured according to panel A. 3. H. Summary of residues in Flu-NP reported to interact with RNA. Flu-NP and all superimposed RNA conformations observed in the TiLV-NP pseudo-C2 and pseudo-C5 structures are transparent. Key interacting residues are displayed as spheres and coloured according to panel A.

**Supplementary Figure 10.**
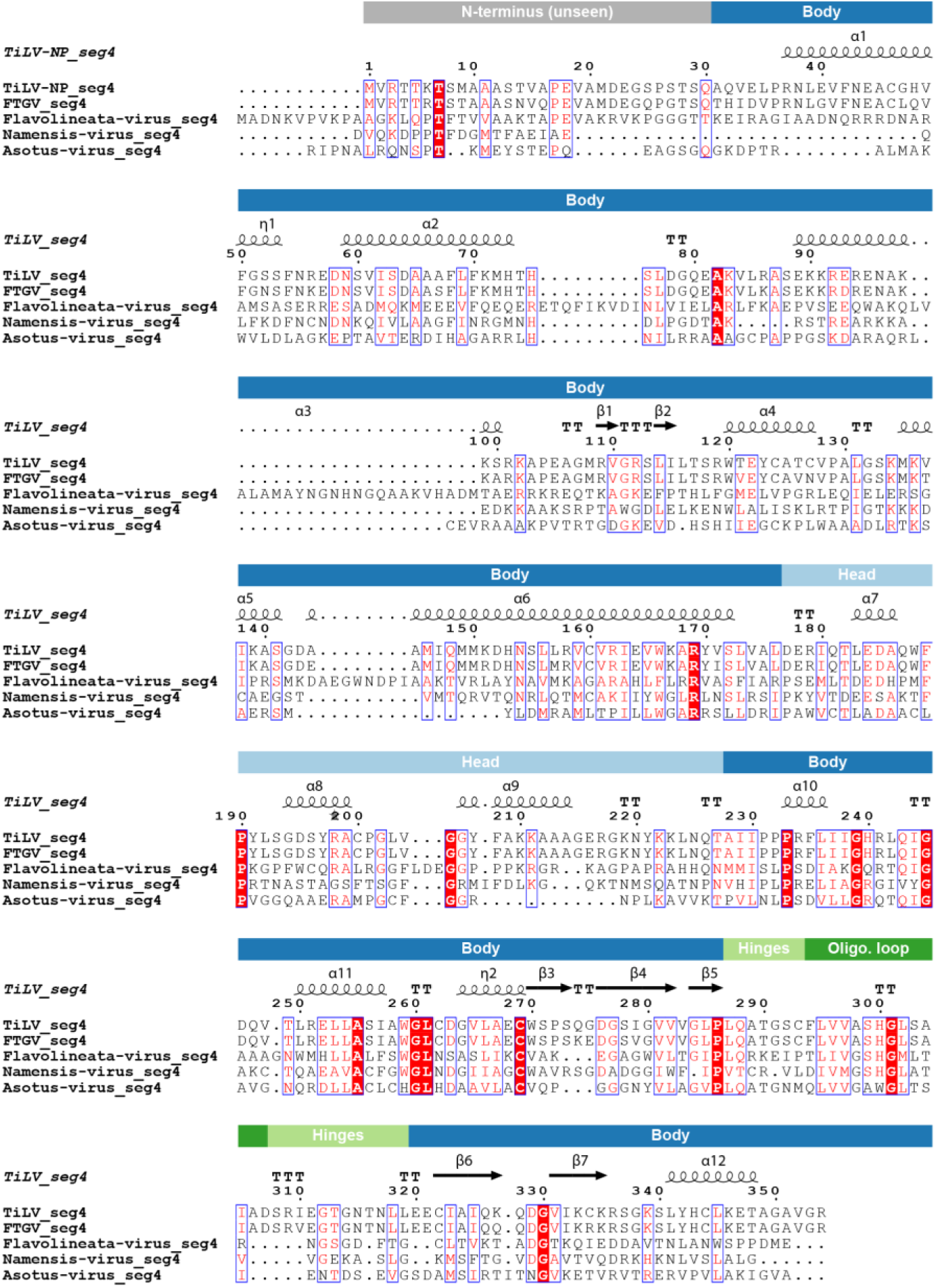
Multiple sequence alignment of TiLV-NP segment 4 with corresponding segment 4 from Fancy tailed guppy virus (*Amnoonviridae* family), Flavolineata, Namensis, and Asotus viruses (not classified). TiLV-NP secondary structures are shown and numbered. Domain positions are indicated. Multiple sequence alignment was performed using MUSCLE (1) with TiLV-NP (segment 4; GenBank: AOE22901.1), Fancy tailed guppy virus (FTGV; segment 4; GenBank: WWU01869.1). Flavolineata, Namensis and Asotus virus putative NP sequences were derived from (2).

**Supplementary Figure 11.**
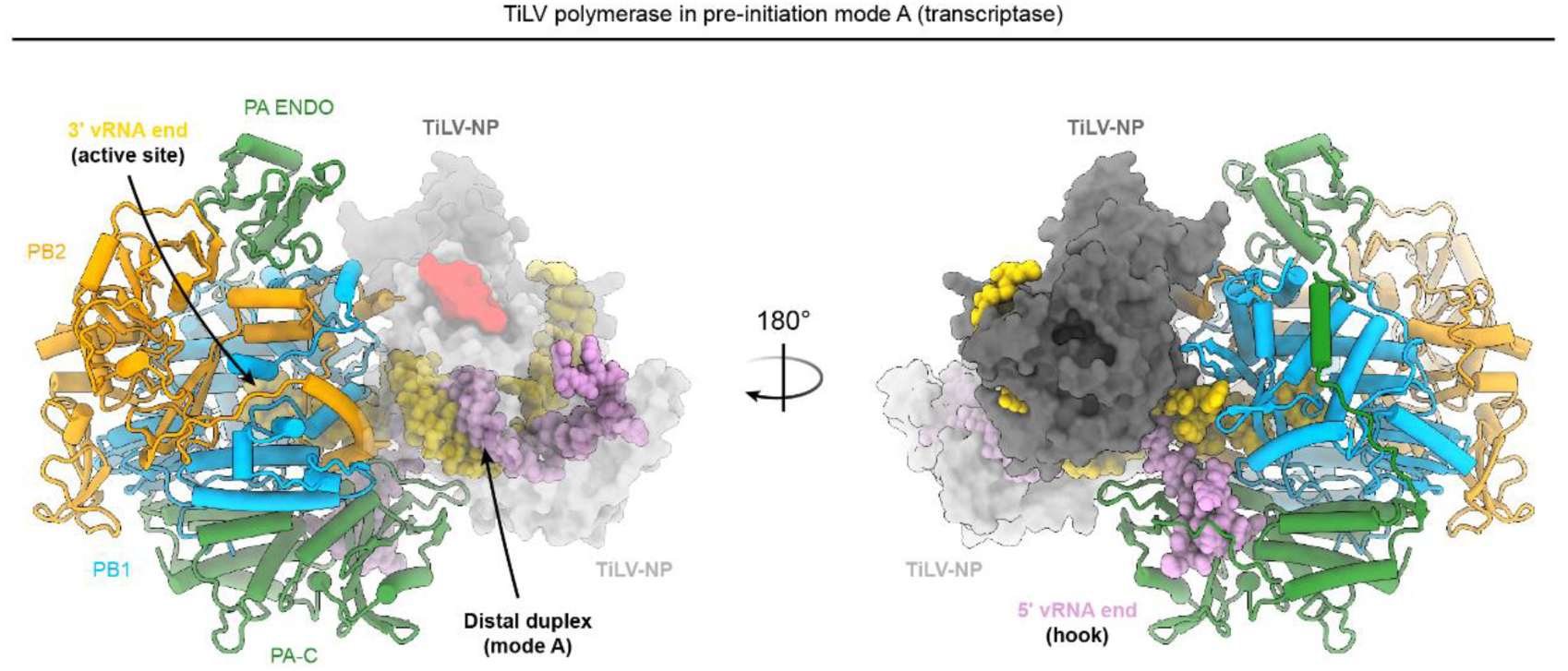
Model of the heterotrimeric TiLV viral polymerase in complex with two TiLV-NPs from the pseudo-C2 tetramer. The A-from RNA observed within the TiLV-NP tetramer (pseudo-C2) is superimposed on the distal duplex of TiLV polymerase (TiLV-Pol) in pre-initiation state mode A with the 3′ end in PB1 active site (PDB: 8PSN). Left: TiLV-NPs are shown as transparent to reveal the distal duplex in mode A. The NP encapsidating the 5′ end (plum) is coloured in light grey with the adjacent NP oligomerization loop in red. The NP encapsidating the 3′ end (gold) is coloured in dark grey. RNA models are displayed as spheres. TiLV-Pol subunits PA, PB1 and PB2 are coloured green, blue, and orange, respectively. Right: 180° rotated view with NPs shown as non-transparent. The 5′ end binds TiLV-Pol as a hook. Few clashes are noted between the NPs and TiLV-Pol, occurring only at flexible loops position.

